# Role of the Arp2/3 Complex in Regulating Mitotic Entry and Progression in *S. pombe*

**DOI:** 10.1101/2025.09.05.674440

**Authors:** Dhanya Kalathil, Erik Flores Reyes, Maitreyi Das

## Abstract

Fission yeast undergoes closed mitosis, wherein chromosome segregation occurs inside an intact nucleus. The microtubule organizing center, Spindle Pole Body (SPB) is duplicated in the G2 phase followed by insertion into a fenestra on the nuclear envelope (NE). The SPB then nucleates spindle microtubules at mitotic onset. Here we show that the branched actin network promotes mitotic entry by enabling NE fenestration. Inhibiting the branched actin nucleator Arp2/3 complex with the drug CK666 fail mitotic entry and the few cells that enter mitosis show significant delays. Upon CK666 inhibition, most proteins are correctly recruited to the SPB at the onset of mitosis however they fail to properly recruit the POLO-like kinase Plo1 and the SPB docking protein Cut11, suggesting a failure in SPB insertion. The start of NE fenestration is marked by the formation of a Sad1 ring, a SUN domain protein followed by a Cut11 ring. Super-resolution structured illumination microscopy indicates that CK666-treated cells show impaired Sad1 or Cut11 rings. Furthermore, these cells do not leak NLS-GFP, a hallmark of fenestration failure. In agreement with the role for branched actin in NE fenestration, CK666 treatment after SPB insertion does not show any mitotic entry defects as observed with preprophase-arrested tubulin mutants. We observed a component of the Arp2/3 complex Arc5, and actin-binding proteins Fimbrin and Coronin, on the nuclear envelope within minutes of mitotic onset. In CK666-treated cells where mitosis is delayed, in the absence of branched actin the SPB aberrantly interacts with the actomyosin ring, preventing its proper dynamics. This results in phenotypes such as the formation of multinucleate structures, asymmetric nuclear division, floppy nuclear membrane, and severe bending of spindle microtubules. Our findings demonstrate a novel role for Arp2/3 complex-dependent branched actin in promoting NE fenestration and SPB insertion during mitosis.

## Introduction

Mitosis is a part of the cell division process where the segregation of duplicated chromosomes occurs equally into two daughter cells [1]. Each step in mitosis, from entry to separation, is controlled by numerous regulators, ensuring high fidelity in this process [2]. The microtubule cytoskeleton, is the main component of the spindle and essential for mitosis [3]. However, emerging research shows that branched actin cytoskeleton, a component previously thought to be unrelated to mitosis, plays a significant role in regulating different aspects of mitosis, further highlighting the complexities of the process. The divergence in the complexities of mitosis, based on nuclear envelope (NE) remodeling, can be seen from unicellular eukaryotes to multicellular organisms where it is classified as open, semi-open, or closed depending on the integrity of the nucleus during mitosis. [4]. Irrespective of the type of mitosis, two daughter cells with an intact nucleus form upon the culmination of division process.

In the mammalian system, open mitosis involves a complete breakdown of the nuclear envelope before mitosis, and it reforms around the nucleus in telophase of the cell cycle[5, 6]. In semi-open mitosis, as in *Caenorhabditis elegans* and syncytial *Drosophila melanogaster* embryos, NE ruptures only at the polar regions [7, 8]. Whereas, closed mitosis occurs in fission yeast, *Schizosaccharomyces pombe*, where mitosis happens inside an intact nucleus. Even though complete nuclear envelope breakdown is absent in *S. pombe*, it relies on NE fenestration to insert the spindle pole body (SPB) into the nuclear envelope at the onset of mitosis. [9].

The spindle pole body in *S. pombe* is a functional equivalent of the centrosome in metazoans, but it differs structurally[10]. The duplication of SPB in *S. pombe* are not well defined. Based on the current knowledge, SPB duplicates in S/G2, starting with the formation of a half bridge, which presumably happens at mitotic exit in the previous cell cycle [11, 12].Thereafter, maturation of SPB occurs in the G2 phase of the cell cycle. SPB has a layered cytoplasmic and electron-dense nuclear compartment connected through fine striations that span the nuclear envelope. The cytoplasmic compartment must be inserted into the nuclear envelope before mitosis, thereby facilitating the formation of spindle and astral microtubules inside the nucleus [13]. Several proteins, including Cut11, Cut12, Kms2, and Brr6, are associated with SPB activation and insertion into the fenestra [14, 15]. Plo1, a mitotic kinase, is recruited to the SPB through its Interaction with SPB proteins such as Pcp1 and Cut12, driving mitotic entry. It has been shown that *cut12.1* forms a monopolar spindle, as it is incapable of activating new SPBs and integrating them into the nuclear envelope. Additionally, the old SPB activation is delayed, and its integration into the nuclear envelope is defective, results in leakage of nucleoplasm into the cytoplasmic compartment [16]. Similarly, Cut11, a transmembrane protein associated with nuclear pores plays a crucial role in SPB insertion into the nuclear envelope. It has been shown that *cut11.1* mutant cells forms monopolar spindle as the newer SPB fail to insert into the NE [17]. Although studies are conducted to decipher the process of SPB insertion, the fundamental process that precedes it, the NE fenestration, has not been fully unraveled. *Fernández-Álvarez A et al* have shown that NE fenestration relies on the centromere and SPB connection via the LINC complex [17]. However, the mechanistic insights of these processes need further investigation.

The cytoskeletal elements consist of actin filaments, microtubules, and intermediate filaments. Microtubules are the major cytoskeletal elements, that play a crucial role in mitosis in addition to their functions in cell polarity, trafficking, and structural support [18].The actin cytoskeleton in fission yeast consists of actin filaments, branched actin, and the actomyosin ring [19]. A key player in the formation of branched actin is the Arp2/3 complex, which acts as a nucleator that binds to a preexisting actin filament, and starts nucleating new branches at an angle of 70° [20, 21]. Actin filaments are associated with structural support, intracellular transport within the cell, and in distributing organelles between the two daughter cells [22]. In addition to their canonical functions actin filaments play a crucial role in mitosis by facilitating spindle positioning and cytokinesis [23]

Branched actin is primarily involved in endocytosis and cell migration [24, 25]. A few studies suggest the association of branched actin with centrosome in metazoans and that opens up new avenues for research. It has been demonstrated that branched actin functions as a diffusion barrier at the centrosome [26] and the interaction of branched actin with the microtubule network is necessary for mitotic spindle formation and chromosome congression [27]. Moreover, perispindle-branched actin is shown to be associated with controlling spindle size [28]. In starfish oocytes, branched actin is required for nuclear envelope breakdown where an Arp2/3 nucleated F-actin shell, forms underneath the nuclear envelope [29]. These studies highlight the multifaceted function of branched actin in cellular physiology.

Here we present a novel finding on the role of branched actin in mitosis. We observe that branched actin inhibition leads to a mitotic entry defect and alterations in spindle dynamics and nuclear fission in *S. pombe*. We find that reorganization of the Sad1 ring, a primary step to NE fenestration, is impaired in cells lacking branched actin. This combined with the fact that cell lacking branched actin do not leak the nuclear marker NLS-GFP indicate a role for branched actin in NE fenestration. We also find the Arp2/3 complex and branched actin binding proteins such as Fimbrin[30] and Arc5-localize to the NE a few minutes before mitosis initiates. Additionally, we find that in the absence of branched actin the nucleus aberrantly interacts with the actomyosin ring leading to abnormal nuclear fission and spindle dynamics during mitotic progression. We demonstrate-that the distorted actomyosin contractile ring observed in CK666-treated cells confers these defects. Overall, our study provides new insights into the role of branched actin in NE fenestration and mitotic progression.

## Results

### Disruption of branched actin leads to mitotic entry defects in *S. pombe*

The small molecule inhibitor CK666 blocks Arp2/3 complex-mediated branched actin nucleation resulting in endocytic and growth defects [30]. Here we report that cells treated with CK666 also show mitotic entry defects. We treated *Bqt4-GFP Sad1-mCherry* expressing cells with 100 μM CK666 and after 5 minutes began assessing the percentage of the mitotic population over a duration of 1 hour. As *S. pombe* cells generally enter mitosis when they reach a length of 14 μm, we only considered the cells in the range of 14-17 μm. Separation of SPB, as evidenced by the splitting of the Sad1 foci, marks mitotic entry. The categorization of cells into different cell cycle stages is accomplished using SPB distance and Bqt4-GFP, an inner nuclear envelope marker. A significant reduction in prophase, metaphase, and early anaphase populations is observed in treated cells compared to DMSO controls (Fig.1A, B). The accumulation of post-anaphase cells in the CK666-treated group indicates effectiveness of the drug, as inhibition of branched actin by CK666 prevents cell separation [31] (Fig.1A and B).

**Figure 1:**
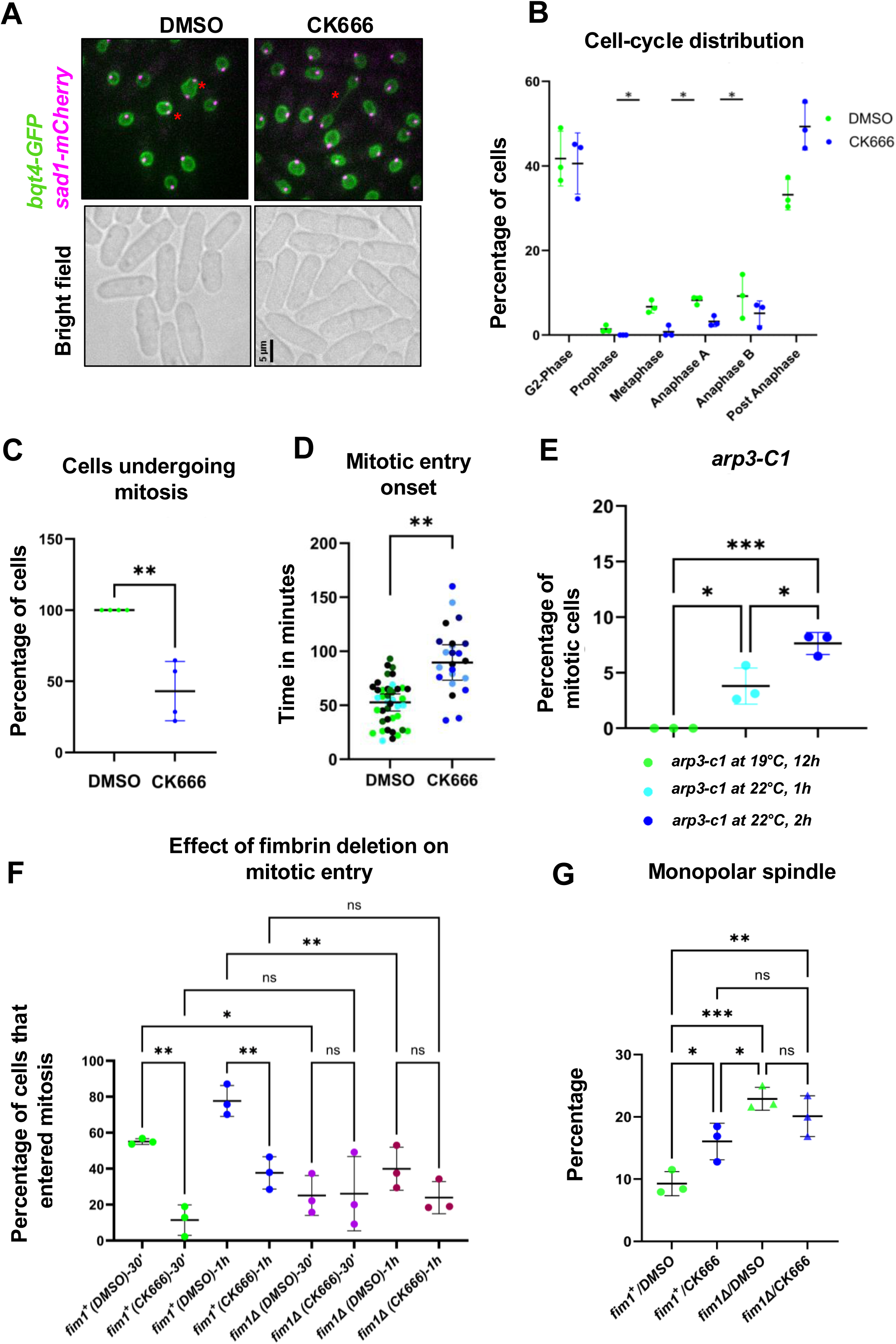
Disruption of branched actin reduces the mitotic cell population. **A.** Sad1-mCherry and Bqt4-GFP expressing cells treated with DMSO or 100μm CK666. Asterisk marks mitotic cells. n ≥ 83, N=3, Scale bar, 5 µm. **B.** Quantification of mitotic population in cells represented in A. **C.** Quantification of mitotic entry in late G2 synchronized *cdc25-22* cells up to 2 hours after release with DMSO or CK666 treatment. n ≥ 15, N=3, student’s t-test. **D.** Quantification of timing of mitotic entry in DMSO and CK666-treated cdc25-22 cells after release from arrest. Error bars show standard deviation, n ≥ 6, N=4, student’s t-test. **E.** Mitotic index of *arp3-C1* mutants at indicated temperatures. n ≥ 55, N=3. one-way ANOVA **F**. Mitotic index after cell cycle release for the indicated times of *cdc25-22 fim1+* and *cdc25-22 fim1ι1* cells treated with DMSO or CK666. n ≥ 58, N=3, one-way ANOVA. G.Monopolar spindle formation in cdc25-22 *fim1+* and *cdc25-22 fim1ι1* cells treated with CK666. n≥63, N=3. One-way ANOVA. *p < 0.05; **p < 0.001, *** p < 0.001, *** p < 0.0001, ns-not significant, mean ± SD.

The Arp2/3 complex causes cell growth arrest [31]. To rule out the possibility that the observed mitotic entry defect is due to inadequate cell size, we used the temperature-sensitive cell cycle mutant *cdc25-22* expressing SPB markers *cdc11-GFP sad1-mCherry*. These cells, at restrictive temperature arrest in late G2 phase and continue to grow to a much larger cell size. After the G2 arrest, cells shifted to the permissive temperature immediately start mitosis. We synchronized *cdc25-22 cdc11-GFP sad1-mCherry* cells in G2 and treated them with CK666/DMSO before shifting them to the permissive temperature (S1A Fig.). Our results show that inhibition of the Arp2/3 complex either fails or delays mitotic entry, with only 43.12% of the cells in the CK666-treated group progressing to mitosis 2 hours after release compared to 100% in control cells (Fig. 1C, S1B). Notably, within the population that did enter mitosis in CK666 treated cells, mitotic entry is significantly delayed, with an average time of 92.91 minutes compared to 51.96 minutes in the controls (Fig. 1D, S1B and Video S1).

To verify that the mitotic defect observed with CK666 treatment is indeed due to inhibition of the Arp2/3 complex, we used the genetic mutant, *arp3-c1* expressing *Atb2-mCherry* labeled microtubules as a marker for mitosis. We find that *arp3-c1* mutant cells grown at a restrictive temperature of 19°C, do not display any mitotic cells as evidenced from the absence of spindle microtubules. However, the mitotic population appears immediately after shifting the cells to a semi-permissive temperature of 22°C (Fig. 1E, S1C). After 1 hour of incubation at 22°C, 3.8 % of the cells contain spindle microtubules, and at 2h the mitotic population increased to 7.6 % (Fig. 1E). Irrespective of the temperature at which the cells are grown, the wild type population (*arp3^+^atb2-mCherry*) contain mitotic cells (19°C-17.67%, 22°C-19.32%), indicating that temperature does not have an influence on mitotic entry (Fig. S1D). These findings show that branched actin disruption hinders mitotic entry.

If the mitotic entry defect is associated explicitly with the absence of the branched actin, we anticipate a similar defect in mutants with disrupted branched actin organization. Fimbrin is an actin bundling protein that aids in organizing actin filaments into higher-order networks [32, 33]. It has been shown that the crosslinking of actin filaments by fimbrin, Fim1 is required for cell division [33]. We assessed mitotic entry in the *cdc25-22 fim1Δ atb2-mcherry* temperature-sensitive strain with and without CK666 treatment. We find that the synchronous *cdc25-22 fim1Δ* cells show significant failure to enter mitosis regardless of DMSO or CK666 treatment. There is no significant difference in the mitotic entry percentage between these groups at 30 minutes and 1h of incubation at permissive temperature irrespective of CK666 (Fig. 1F, S1E), whereas *cdc25-22 fim1+* cells, only show defects in mitotic entry upon CK666 treatment (Fig. 1F). These results provide further evidence that branched actin has a role in mitotic entry. In addition, we find enhanced monopolar spindle formation in *fim1ι1* compared to *fim1^+^*cells which have undergone CK666 treatment (Fig. 1G, S1E).

Cellular stress leads to a mitotic entry defect by sequestering the Cdc25 protein in the cytoplasm[34, 35]. To rule out that the mitotic defects in CK666 cells were due to the stress response we analyzed the nucleo-cytoplasmic ratio of Cdc25mNG after treatment with CK666. Cells treated with the actin inhibitor latrunculin A (LatA) and with high salt concentration (KCl) are known to sequester Cdc25-mNG in the cytoplasm. We observed that unlike LatA and KCl, CK666 treatment did not sequester Cdc25-mNG in the cytoplasm, instead, it was observed in the nucleus, indicating the absence of a stress response (Fig. S2A and S2B).

### Spindle Pole Body insertion is impaired upon branched actin disruption

Given the mitotic entry defects observed upon branched actin disruption, we examined the molecular players involved in this process. The Polo kinase, Plo1 localizes to the spindle pole body (SPB) immediately before the onset of mitosis and signals mitotic commitment [36]. We monitored Plo1-GFP recruitment to the SPB in synchronized *cdc25-22 plo1-GFP sad1-mCherry* cells treated with CK666. The Plo1-GFP signal at the spindle pole body gradually increases through the late G2 phase of the cell cycle, approximately 8-10 minutes before the onset of mitosis as marked by SPB separation (Fig. 2A). However, in CK666-treated cells we find abnormal patterns of Plo1-GFP recruitment to the SPB. One group of cells failed to recruit Plo1-GFP to the SPB (Fig..2A, B, CK666-Phenotype 1 and Video S2). We find that 73.54% and 49.03% of the cells are devoid of Plo1 at the SPB at 30 minutes and 1h after the release to the permissive temperature of 25°C, compared to DMSO-treated controls (30min-17.17%, 1h-1.010%) (Fig. 2A and B). Another subset of cells shows delayed Plo1-GFP recruitment to the SPB post shifting to permissive temperature (Fig. 2A, CK666-Phenotype 2 and Video S2). In a third subset of cells Plo1-GFP signal appears immediately after the cells are released into permissive temperature but mitotic onset is delayed and Pol1-GFP continues to accumulate at the SPB (Fig. 2A, CK666-Phenotype 3 and Video S2).

**Figure 2:**
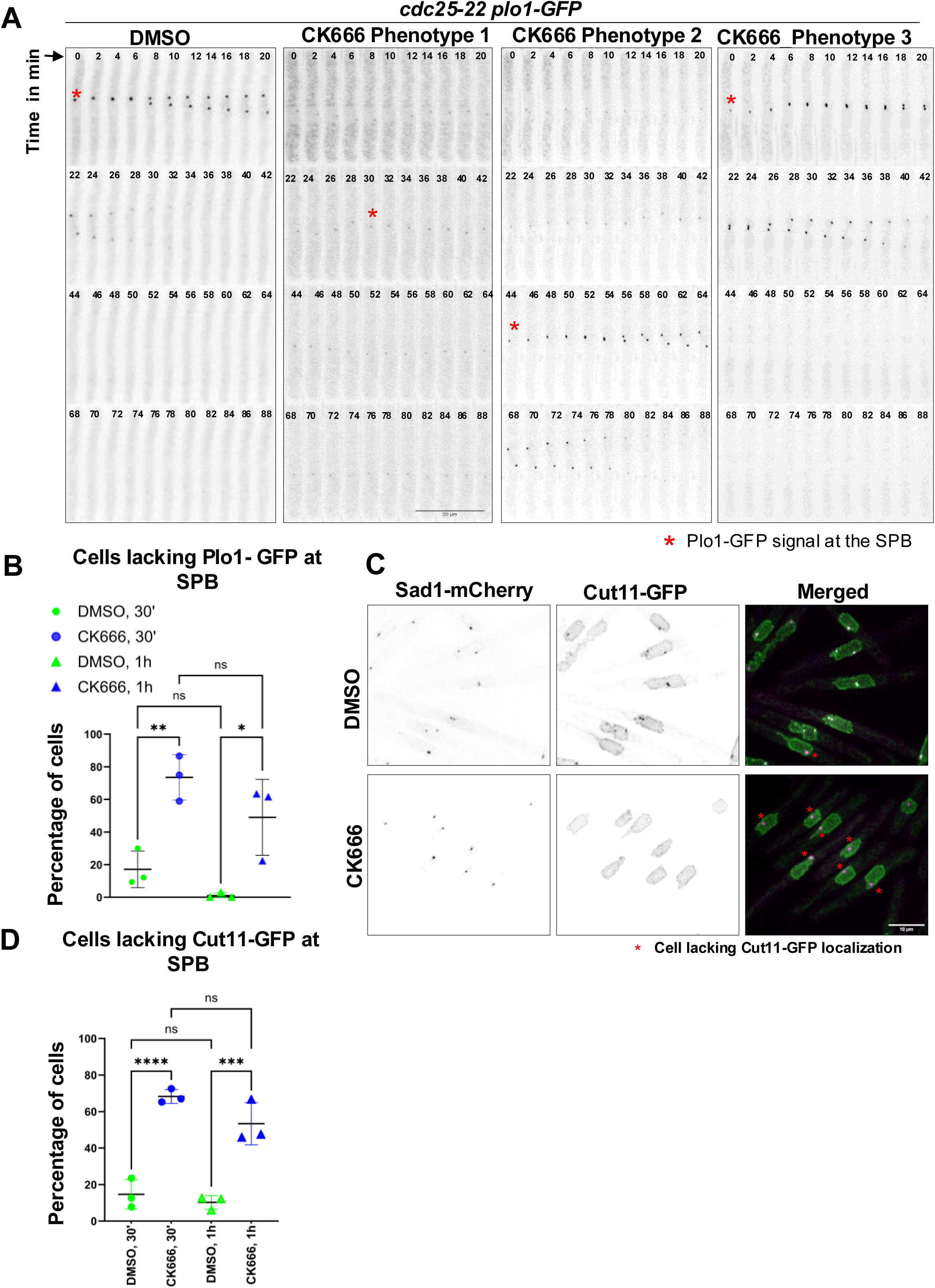
Loss of branched actin impairs Plo1 and Cut11 recruitment to the SPB. **A.** Plo1-GFP localization in DMSO or CK666-treated synchronous *cdc25-22* cells released into the cell cycle. Plo1-GFP localization at the SPB is failed in CK666 phenotype 1, delayed in phenotype 2 and recruited earlier but the mitotic onset is in phenotype 3. Scale bar-20 µm. **B.** Quantification of cells lacking Plo1-GFP at the SPB in *cdc25-22* cells after cell cycle release with DMSO or CK666 treatment. n ≥ 30, N=3. **C.** Cut11-GFP recruitment to the SPB in synchronized cdc25-22 cells released into the cell cycle after DMSO or CK666-treatment. Sad1-mCherry marks the SPB. Red asterisk marks cells lacking Cut11 at the SPB. Scale bar, 10 µm. **D.** Quantification of cells failing to recruit Cut11-GFP to the SPB of cells represented in C. n ≥ 66, N=3.One-way ANOVA. *p < 0.05; **p < 0.01, *** p < 0.001, *** p < 0.0001, **** p < 0.00001 ns-not significant, mean ± SD.

Cut11 is a spindle pole body docking protein required for SPB’s insertion into the nuclear envelope. In interphase Cut11 is observed throughout the nuclear envelope and is recruited to the SPB at the time of mitotic entry where it remains until early anaphase [37]. The synchronized *cdc25-22 sad1-mCherry cut11-GFP* cells treated with CK666, fail to recruit Cut11 to the SPB (Fig. 2C). At 30 minutes and 1hour after the release to permissive temperature, 68.25% and 53.35% of cells, respectively, are without Cut11, compared to 14.66% and 10.27% in DMSO-treated cells (Fig. 2D).

It is possible that the Plo1 and Cut11 recruitment defects observed in CK666 cells is due to improper recruitment of proteins to the spindle pole body after the previous cell cycle. To test this, we assessed the intensity of Cdc11, a protein present at the SPB throughout the cell cycle, and Pcp1, a protein required for Plo1 and γ-tubulin ring complex recruitment, in *cdc25-22* synchronized cells treated with CK666. Both Cdc11 and Pcp1 levels gradually increase in the SPB from the G1/s to G2 to M phase of the cell cycle[38]. If CK666 treatment impairs the assembly and recruitment of spindle pole body proteins the Cdc11 and Pcp1 levels would be decreased in these cells. Instead, we find that both Cdc11-GFP and Pcp1-GFP intensity remains unchanged -in CK666 treated cells compared to DMSO controls at 0, 10, and 20 minutes after shifting the cells to a permissive temperature (Fig. S3A and B). Moreover, we did not observe any change in the intensity profile of Sad1-mCherry (Fig. S3C). This suggests that the proteins required for proper SPB assembly are correctly recruited to the SPB and CK666 likely impacts events after this process resulting in mitotic entry defects.

### Nuclear envelope fenestration is impaired in branched actin disrupted cells

Following SPB protein recruitment and duplication, the new SPB is inserted into a fenestra in the NE. To determine whether the fenestration process is impaired, ensuing branched actin inhibition, we sought to investigate the status of Sad1 ring formation using structured illumination microscopy of fixed, synchronized cells over a period of 30 minutes after release from late G2 phase arrest. Sad1 is a component of the LINC complex, and its redistribution into a ring-like structure marks the beginning of the fenestration process [39, 40]. We find that either the Sad1 ring formation is completely absent or improperly formed in CK666-treated cells compared to controls (Fig. 3Ai). Sad1 ring is formed in only 14.07% of the cells compared to 38.35% in DMSO-treated cells (Fig. 3B). In addition to the Sad1 ring, we find that Cut11-GFP also forms a ring that colocalizes with the Sad1 ring in the control cells. While we see some cells with only a Sad1-mCherry ring, we never see a Cut11-GFP ring by itself (Fig. 3Aii). This suggests that the Cut11-GFP ring forms after the Sad1 ring. We find that only 3.427% of the cells in CK666-treated cells contain both sad1-mCherry and cut11-GFP rings compared to 21.46% of the rings in the controls (Fig. 3C).

**Figure 3:**
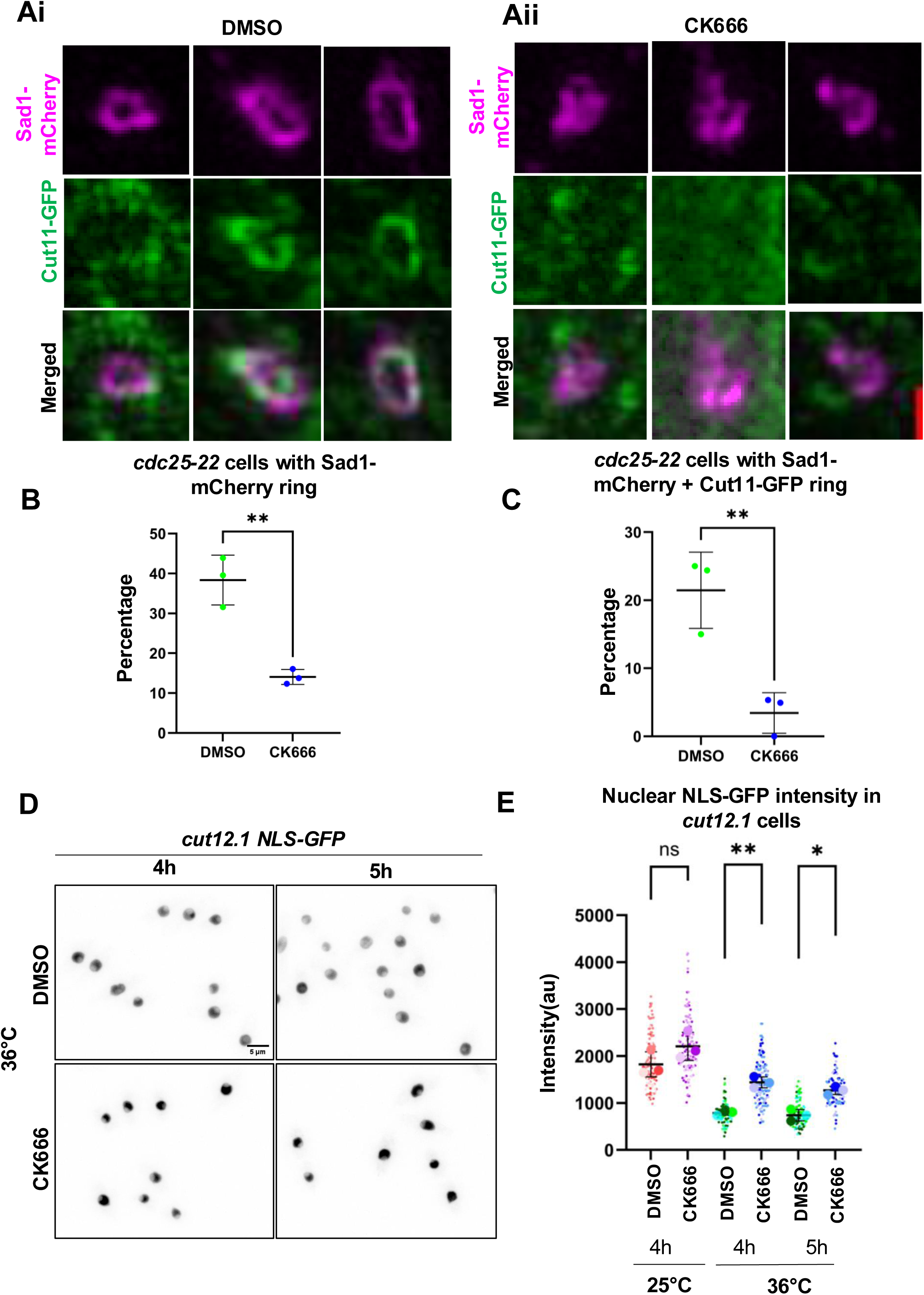
NE fenestration is compromised in cells lacking branched actin. **A.** Redistribution of Sad1-mCherry and Cut11-GFP in synchronized cdc25-22 released into the cell cycle for 30 mins after DMSO or CK666 treatment. Scale bar-0.5 μm. **B.** Quantification of cells displaying Sad1-mCherry rings in cells described in A. **C.** Quantification of cells displaying both Sad1-mCherry and Cut11-GFP ring at the SPB in the cells described in A. n ≥ 40, N=3, mean ± SD student’s t-test. **D.** Representative images display accumulation of NLS GFP in the nucleus in DMSO or CK666 treated *cut12.1 NLS-GFP* strain grown at 36°C for 4 and 5 hours. Scale bar-5 µm. **E.** Quantification of NLS-GFP intensity in the nucleus in *cut12.1* cell grown at 25°C and 36°C, n ≥ 36, N=3, mean ± SD, One-way ANOVA *p < 0.05; **p < 0.001.

The absence of Sad1 ring formation implies the absence of fenestra formation. To further confirm that branched actin disruption hinders the fenestration process, we used the *cut12.1* strain expressing NLS-GFP. The *cut12.1* mutant restrictive conditions forms a fenestra but fails to embed one of the duplicated SPBs into the nuclear envelope and thus leaks NLS-GFP [16]. We treated *cut12+* and *cut12.1* cells with CK666/DMSO and imaged at regular intervals for a period of 6hrs at restrictive temperature to monitor NLS-GFP signal in the nucleus. If branched actin inhibition leads to impairment of the NE fenestration, we expect to see the accumulation of NLS-GFP in the nucleus even in the *cut12.1* mutant [17, 39]. Indeed, we find that in CK666-treated cells, NLS-GFP in the nucleus accumulates compared to the controls (Fig. 3D and E), supporting a role of branched actin in NE fenestration.

It has been reported that SPB insertion defects lead to a delay in mitotic onset after depolymerization of the cytoplasmic microtubules [13, 15]. We asked if such a delay also occurred in the fraction of CK666-treated cells that were able to enter mitosis. We observe a delay in mitotic onset after complete depolymerization of the microtubules in cells treated with CK666 (Fig. S4), however, the data while trending is statistically insignificant. We also saw a similar trend in *fim1ι1* mutants that were able to enter mitosis, with or without CK666 treatment (Fig. S4).

### Loss of branched actin do not block mitotic entry in cells released from pre-prophase arrest

At the onset of mitosis, the duplicated SPB is inserted into the nuclear envelope, start nucleating spindle microtubules and the SPBs are separated. The tubulin mutant *nda3-KM311* is cold-sensitive and arrests in preprophase of the cell cycle under restrictive conditions as it fails microtubule nucleation [41]. Cells arrested in pre-prophase complete fenestration and SPB insertion and upon release to the permissive temperature enter mitosis successfully. Thus, these mutant cells should be resistant to CK666 treatment after arrest. The arrested cells when released after treatment should progress to mitosis as fenestration and insertion is already completed. The *nda3-KM311* mutant cells expressing *sad1-mCherry* as a SPB marker were arrested at pre-prophase of the cell cycle, followed by drug treatment and released to permissive temperature and imaged. After 15 minutes of incubation with the drug at 18°C, the cells were shifted to permissive temperature (34°C) and imaged over time (Fig. 4A). We find that, irrespective of the DMSO or CK666 treatment, the cells progress to mitosis, as evidenced by the 75.18% and 68.93% of the mitotic population in the control and test groups, respectively, within 5 minutes after the shift. Similarly, after 15 minutes of incubation at 34°C, a peak in the mitotic population is observed, with 93.32% and 95.13% in the control and test groups, respectively (Fig. 4B and 4C). These results imply that the mitotic entry defects occur before microtubule nucleation.

**Figure 4:**
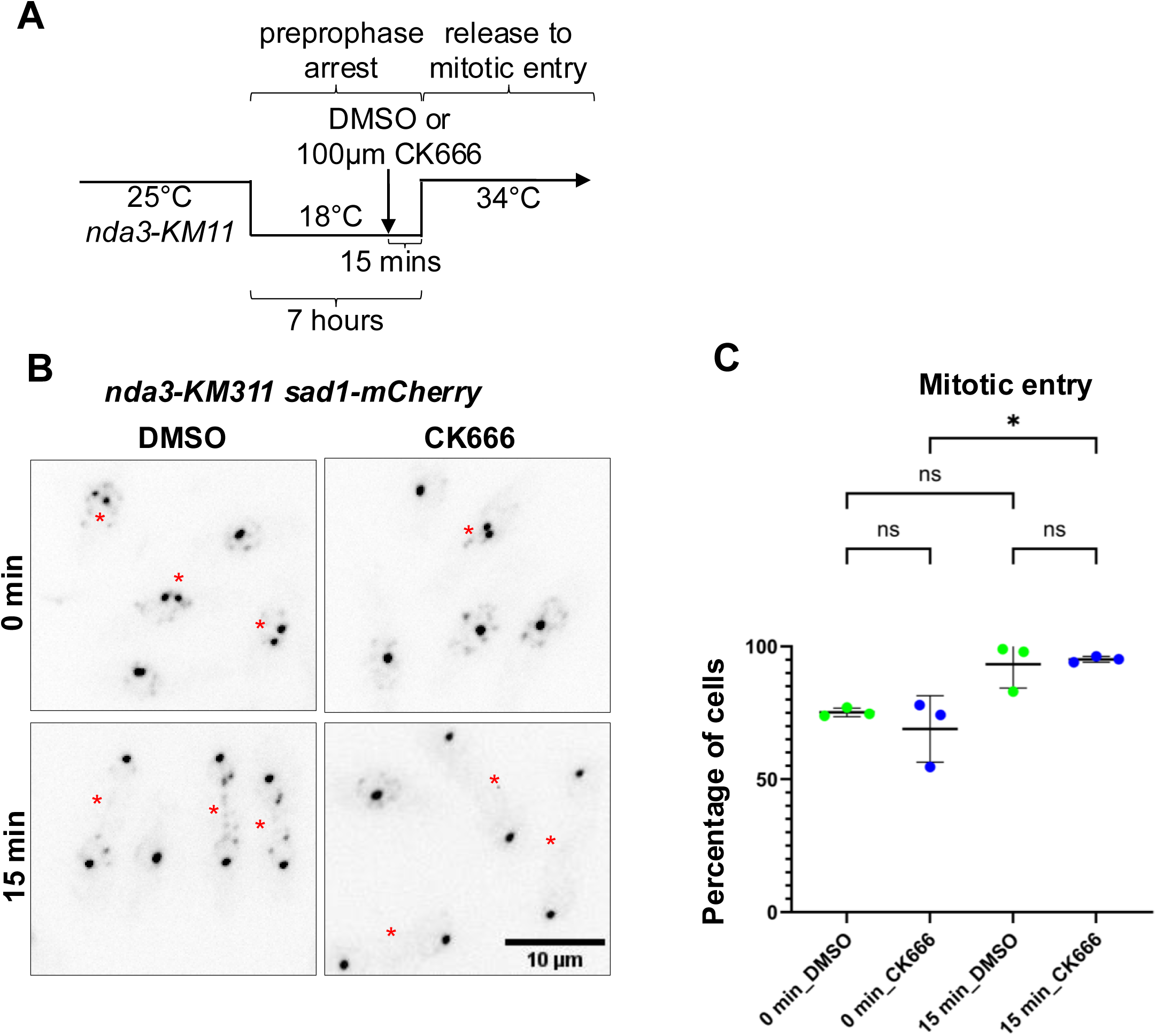
Loss of branched actin after preprophase arrest does not block mitosis when released. **A.** Mitotic progression in *nda3-KM311* cells after release from preprophase arrest in DMSO or CK666 treated cells. Sad1-mCherry marks SPB. Red asterisks represent cells in mitosis. Scale bar-10 µm. **B.** Quantification of the percentage of cells that progress to mitosis at respective time intervals after the release from prophase arrest. n≥50, N=3, mean ± SD, One-way ANOVA, *p < 0.05, ns-not significant.

### Arp2/3 complex associated molecules localize to the nuclear envelope and SPB

Our data suggests that branched actin disruption impairs NE fenestration, thus blocking SPB insertion into the nuclear envelope. We asked whether branched actin is directly involved in these processes. A few studies in metazoans indicated the presence of branched actin at the centrosome and also near the NE [42, 43]. It has been shown that the NE breakdown in star fish oocytes necessitates the formation of an F-actin shell under the nuclear envelope [43]. Based on our observations and the existing knowledge, we speculate that branched actin may directly regulate the NE fenestration. As NE fenestration is a dynamic process, it is challenging to verify this possibility. To better understand this, we examined the presence of actin-related molecules on or near the nuclear envelope and SPB using super-resolution live-cell microscopy. Arc5, Fimbrin, and Coronin-expressing cells were used to elucidate this. Intriguingly, we find that all of these molecules transiently come in contact with either the nuclear envelope (Bqt4-GFP or Bqt4-mCherry) or the SPB (Ppc89-mCherry) (Fig. 5Ai, ii and iii, Table S1-4). We considered up to 15 minutes before SPB separation in our analysis. In the case of Bqt4-GFP expressing cells, as there is no SPB marker to time the mitotic entry, the onset of anaphase (start of nucleus elongation) is considered the 0-time point. The cells initiate anaphase ∼8 minutes after the SPB separation; therefore, we followed the cells up to 18 minutes prior to anaphase onset. This includes 10 minutes before mitotic onset. Although we observe a prominent association of these molecules at the SPB and NE during this timeframe, predicting a trend is challenging as it lacks precise consistency, making the quantification difficult. Due to this, we represent the localization data in a table format. We find that Fim1-GFP colocalizes with Ppc89-mCherry in mitotic cells transiently within a time window of -10 to 0 minutes before mitotic onset (Fig. 5Bi, Table S1 and Video S3). Fim1-GFP also associated with the NE labeled with Bqt4-GFP about -18 to -8 minutes before anaphase onset (Fig. 5Bii, Table S2 and Video S4). We also observed the Arp2/3 complex component Arc5-mCherry associates with the Bqt4-GFP labeled NE -28 to -8 minutes before anaphase onset (Fig. 5Biii, Table S3 and Video S5). Finally, we also saw the actin binding protein coronin Crn1-GFP localizing with the SPB marker Sad1-mCherry -12 to -1 minute before mitotic onset (Table S4). The localization data show that these molecules come into contact with the NE and SPB at the onset of mitosis and are highly dynamic strongly supporting the role of branched actin in NE fenestration, but warrants further investigations.

**Figure 5:**
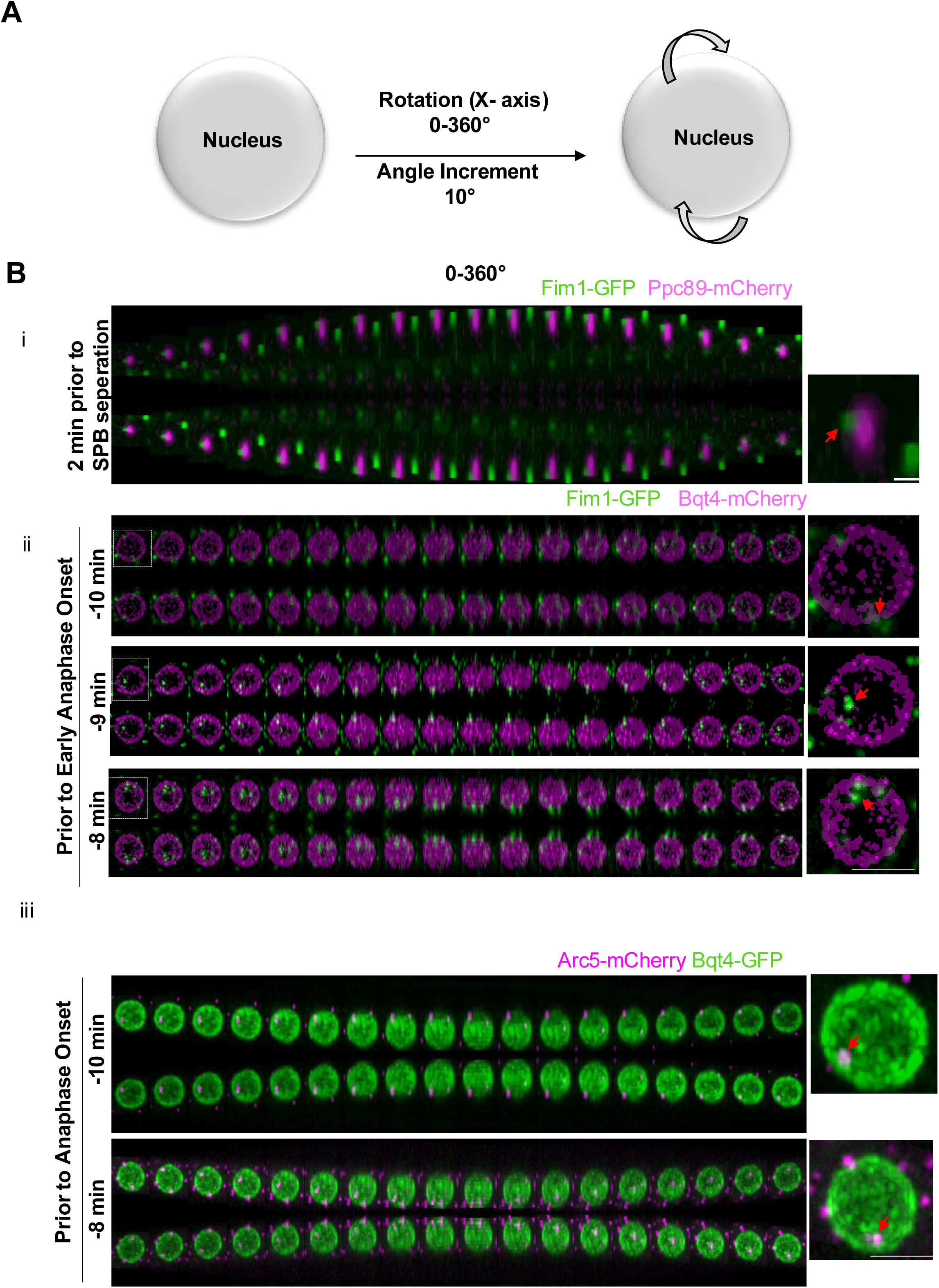
Localization pattern of Arp2/3 related molecules at the SPB and nuclear surface. **A.** Schematic describing angular view (0-360°) of 3D reconstructed nucleus below. **B.** Super-resolution microscopy showing (i) Fim1-GFP localizes adjacent to Ppc89-mCherry 2 minutes prior to onset of mitosis, Scale bar-0.5 μm (ii) Fim1-GFP localizes to Bqt4-mCherry labeled NE -10, -9-, and -8 minutes prior onset of anaphase (iii) Arc5-mCherry localizes to Bqt4-GFP labeled NE - 10 and -8-minutes prior onset of anaphase. Scale Bar-2 μm

### Branched actin disruption leads to defects in nuclear fission and spindle microtubule dynamics

In the fraction of CK666-treated cells where mitosis is not entirely blocked but delayed (Fig.1D, S1B), the cells progress through mitosis, with a series of abnormalities in nuclear fission. Late G2 phase synchronized *cdc25-22 bqt4-GFP* cells were treated with CK666, followed by live-cell imaging at the permissive temperature. Bqt4-GFP and Atb2-mCherry were used as markers for the nuclear envelope and spindle microtubules, respectively. We find a floppy and disorganized nuclear envelope, often forming trinuclear structures, in CK666-treated cells (Fig. 6A and Video S6). Tri-nucleated cells are present in 66.66% of CK666-treated cells compared to the control (8.460%) (Fig. 6A, B, and C). We noted that in cells with defects in nuclear fission, the spindle elongation occurs only in one half of the cell (Fig. 6A). To further investigate this, we examined spindle microtuble dynamics in CK666-treated *cdc25-22 atb2-mCherry* cells still able to enter mitosis. In these cells, the spindle microtubule is severely bent in 59.44% of the cells compared to the DMSO control (0.8037%) when they approach late anaphase of the cell cycle (Fig. 6D, E and Video S7). However, in some instances, the severely bent pattern is rectified to a linear morphology as the nuclear fission process nears its completion.

**Figure 6:**
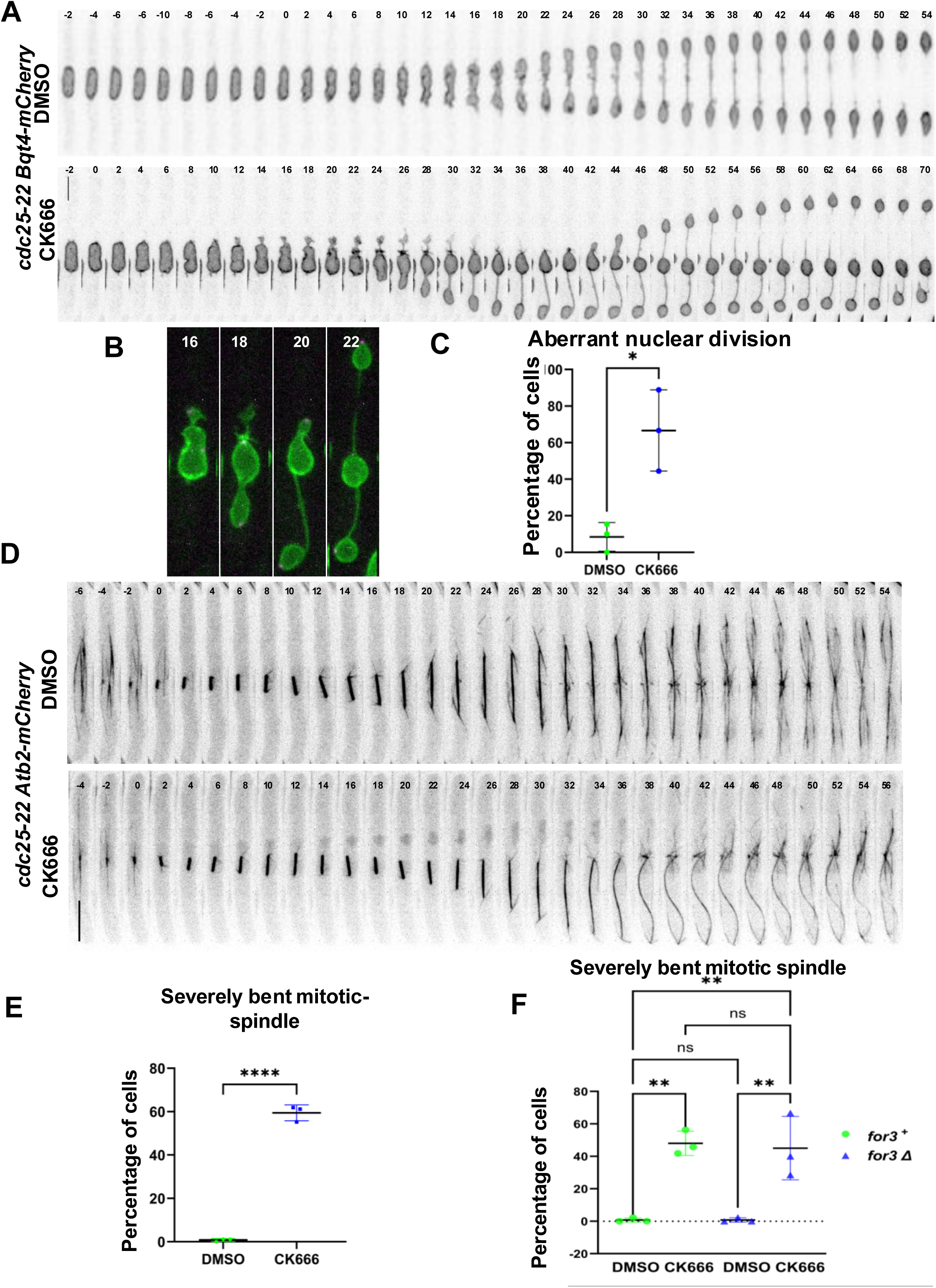
Loss of branched actin results in abnormal nuclear fission. **A.** Nuclear fission in DMSO or CK666-treated synchronized *cdc25-22* cells post cell cycle release. NE is labeled with Bqt4-GFP. Scale bar-10 µm. **B.** Images highlighting nuclear fission defects and multinuclear structures at selected time frames shown in A. **C.** Quantification of aberrant nuclear division in DMSO or CK666-treated cells described in A. n≥6, N=3. Student’s t test. **D.** Atb2-mCherry labeled spindle microtubule dynamics in DMSO or CK666-treated synchronized *cdc25-22* cells post cell cycle release. Severe bending of microtubules is seen in CK666-treated cells. Scale bar-10 µm. **E.** Quantification of severe microtubule bending in DMSO or CK666-treated *cdc25-22 atb2-mCherry* cells. Scale bar-10 μm, n≥50, N=3, n=10, Student’s t test. **F.** Quantification of spindle microtubule bending defects in *for3^+^* cdc*25-22* and *for3Δ* cdc*25-22* cells treated with DMSO or CK666. n≤70, N=3. mean ± SD, one-way ANOVA. *p < 0.05; **p < 0.001. ns-not significant.

To maintain actin homeostasis, in the absence of branched actin, cells produce more linear actin filaments [44]. We asked if excessive linear actin filaments in the cytoplasm interfere with nuclear division. Linear actin filaments in vegetative cells are nucleated by the Formin, For3 [45–47]. In *for3ι1* mutants linear actin filaments are absent. We treated late G2 synchronized *for3Δ cdc25-22 atb2-mCherry* cells with CK666/DMSO and then shifted to permissive temperature followed by live-cell imaging. We find that 45.08% of the *for3Δ* strain treated with CK666 has severely bent microtubules, compared to DMSO-treated cells (0.7937%) (Fig. 6F, S5A). A similar trend (48.03% cells with bent microtubule) is observed with CK666 treatment in a *for3^+^* background. This indicates that nuclear division and microtubule bending defects is not due to excessive linear actin filaments in CK666 cells.

A careful analysis of nuclear division dynamics in these cells appear to show that a part of the nucleus is caught at the cell middle. In these dividing cells the actomyosin ring is present at the cell middle, thus we asked if this structure impedes nuclear division in CK666-treated cells. To address this, we disrupted the contractile ring using a mild treatment of the actin polymerization inhibitor, Latrunculin A[48]. Briefly, synchronized *cdc25-22 bqt4-GFP atb2-mCherry* cells underwent combinatorial treatment with CK666 and LatA to disrupt all the actin structures in the cells, followed by release to permissive temperature, and live-cell imaging. We find that with the combined LatA and CK6666 treatment, which disrupts both actin filaments and branched actin, the cells continue to show mitotic entry defects but do not show the severely bent microtubule phenotype seen with CK666 treatment (Fig. 7A & B). Consistent with our previous results, 55.30 % of the CK-666 treated cells showed bending phenotype compared to control (1.688%). Notably, combinatorial treatment rescued the severely bent phenotype, with only 3.512% of the cells displaying this defect. (Fig. 7B). These results suggest that the nuclear division and microtubule bending defects occur due to aberrant interaction between the nucleus and the actomyosin ring.

**Figure 7:**
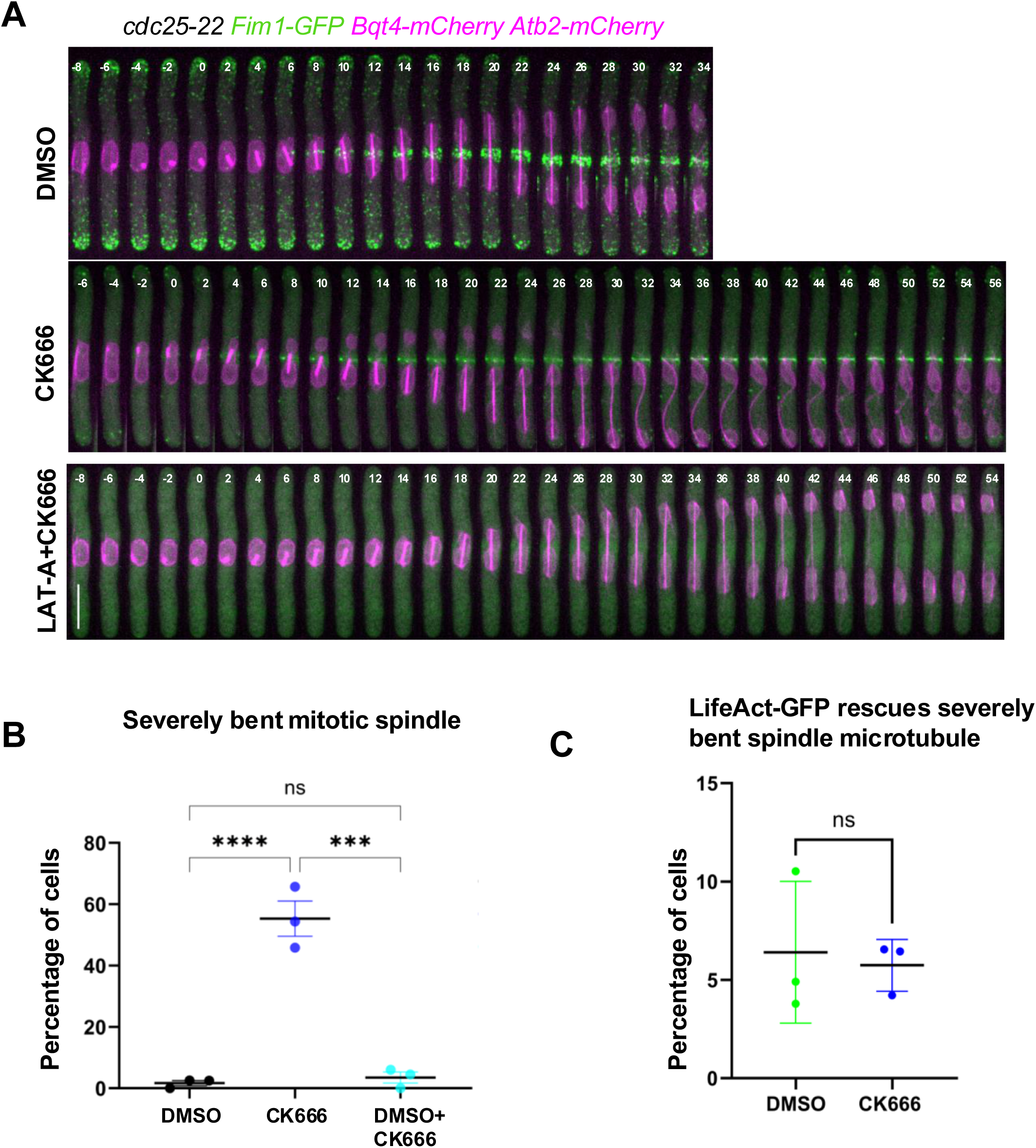
Spindle microtubule bending defects require an intact actomyosin ring. **A.** Spindle microtubule dynamics in DMSO, CK666, and CK666+LATA treated *cdc25-22 fim-GFP bqt4-mCherry atb2-mCherry* cells. Scale bar: 10 μm (B). Quantification of spindle microtubule bending defects in cells shown in A. n ≥ 24, N=3, mean ± SD, Ordinary one-way ANOVA. **p < 0.001, *** p < 0.001, ns-not significant. **C.** Quantification of spindle microtubule bending defect in *cdc25-22* cells expressing LifeAct-GFP treated with DMSO or CK666. n≥19, N=3, mean ± SD, Student’s t-test, *p < 0.05; *** p < 0.001, ns-not significant.

To further assess this, we analyzed *cdc25-22* cells expressing *LifeAct-GFP* as a marker for F-actin. We find that CK666-treated, synchronized LifeAct-GFP cells released to a permissive temperature do not exhibit defects in either nuclear or spindle microtubule dynamics but continued to show mitotic entry defects (Fig. 7C, S5B and C). The LifeAct-GFP probe functions by binding a hydrophobic site on the actin subunits. This site also interacts with other actin binding proteins that LifeAct-GFP can outcompete [49]. It is possible that in the CK666 treated cells LifeAct mask the actin molecules on the ring thus preventing interaction with the nucleus. This further supports our hypothesis that in the absence of branched actin the nucleus interacts with the actin on the contractile ring and this prevents proper nuclear division resulting in microtubule bending.

As we have seen that the disruption of the contractile ring rescued the severely bent microtubule phenotype of CK666-treated cells, we sought to determine how the contractile ring contributes to the nuclear fission defects and severely bent microtubule phenotype. To investigate this, synchronized *cdc25-22 cdc11-GFP rlc1-tdTomato* cells were treated with CK666/DMSO, shifted to permissive temperature and then imaged over time. Rlc1-tdTomato and Cdc11-GFP mark the contractile ring and SPB, respectively. In DMSO treated cells, the nucleus elongates during mitosis and the SPB moves to either side of the cell medial region. At the onset of mitosis, both the SPBs are visible at the plane of the actomyosin ring (Fig. 8A, B and Video S8). As mitosis progresses the SPBs move away from the ring, and towards the two cell ends. In CK666 treated cells with nuclear elongation defects, at least one SPB appears to linger at the cell medial region Fig. 8A, B and Video S8). We find that with mitotic progression, one SPB moves away from the plane of the ring while the other appears to be caught with the ring. Moreover, 3D-reconstruction of the actomyosin ring shows components of the Rlc1-tdTomato labeled ring being pulled by the Cdc11-GFP marked SPB. (Fig. 8A). We observed this aberrant interaction between the actin ring and the SPB in about 75% of the cells that were able to enter mitosis upon CK666 treatment (Fig. 8C). These observations suggest that the SPB at the nucleus interacts with actin in the contractile ring in the absence of branched actin, thereby restricting proper nuclear fission, and resulting in severe spindle bending defects.

**Figure 8:**
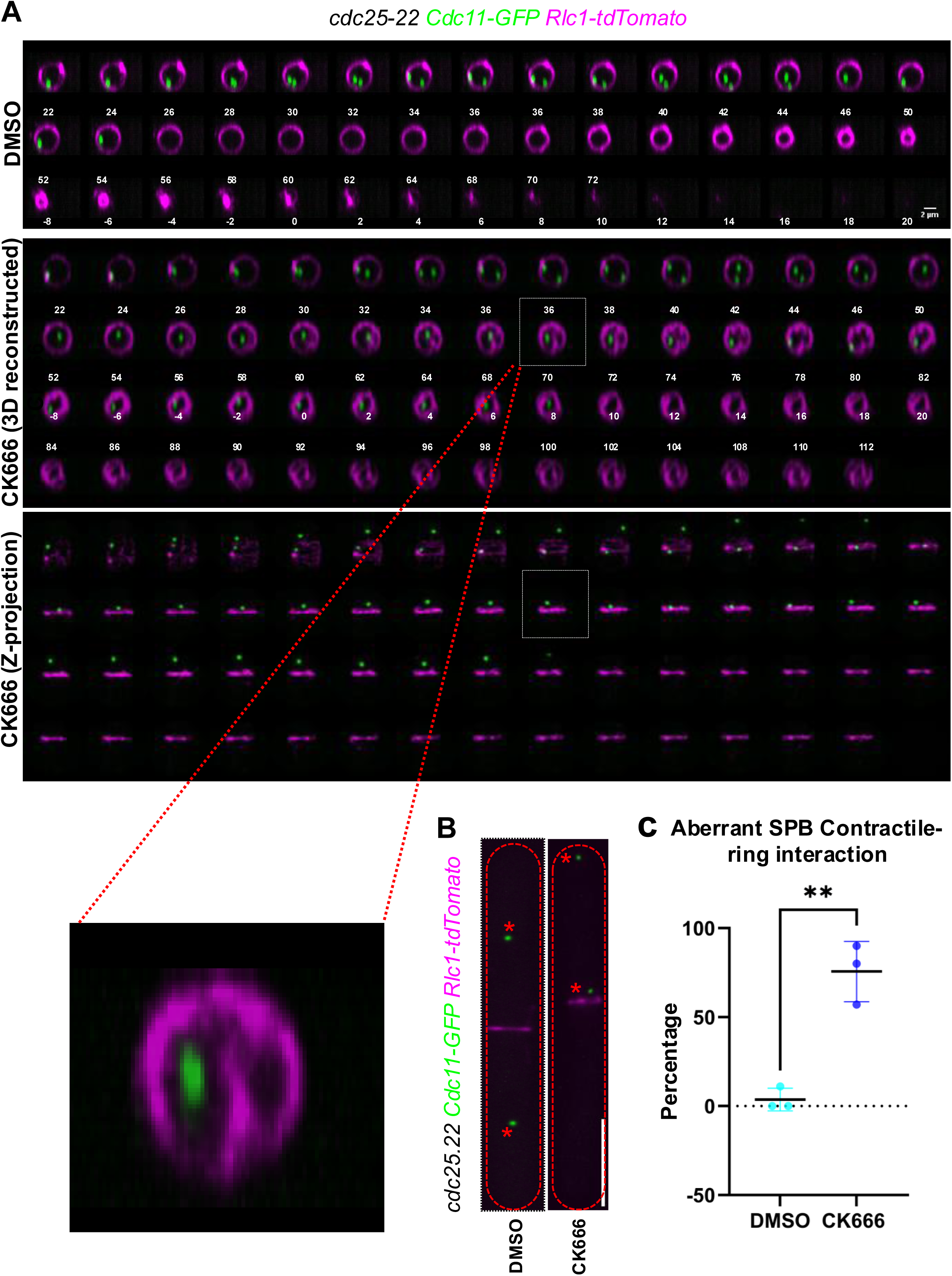
In the absence of branched actin, the SPB aberrantly interacts with the actomyosin ring. **A.** 3D-reconstruction over time of SPB labeled with cdc11-GFP and the actomyosin ring labeled with Rlc1-tdTomato in DMSO or CK666-treated *cdc25-22* cells. Enlarged image of a SPB aberrantly positioned at the plane of the actomyosin ring and inward protrusion of the ring. Scale bar-2μm. **B.** Representative cells shown in A. Scale bar-10 μm. **C.** Quantification of aberrant SPB and actomyosin ring interaction in DMSO or CK666-treated cells. n≥7 cells, N=3. p value, * <0.05, Student’s t test.

## Discussion

The SPB in *S. pombe* acts as a microtubule organizing center, giving rise to cytoplasmic microtubules during cell cycle interphase. During mitosis, it nucleates spindle microtubules [50]. The spatiotemporal coordination of these dual roles involves SPB’s insertion-extrusion cycle into and out of the nuclear envelope, respectively [13]. In mitosis, after the SPB is successfully inserted, the nucleoplasmic phase gives rise to spindle microtubules. As *S. pombe* undergoes closed mitosis, this process necessitates NE fenestration for SPB insertion [14]. The mechanistic details of NE fenestration are still elusive. We find that branched actin promotes NE fenestration and in cells lacking branched actin mitotic entry is disrupted.

Here we show a mitotic entry delay or failure upon either chemical inhibition of branched actin nucleation or with genetic mutants. Our studies further inform that the defects lie between late G2 and pre-prophase of the cell cycle, as based on our observation with *cdc25-22* and tubulin mutants. The defects in mitotic entry could be attributed to improper recruitment or assembly of SPB proteins. After each cell cycle, the SPB recruits specific proteins to enable its duplication and assembly and this is then inserted into the NE just before mitotic entry [38]. We did not see any defects in the levels of SPB proteins such as Pcp1, Cdc11 and Sad1 suggesting that CK666 impacts the SPB after its maturation and duplication.

The Polo-like kinase1 (Plo1), a mitotic regulator, is associated with cell-cycle control, Septation Initiation Network signaling, and stress-response MAP kinase pathways [36, 51]. We find that Plo1 recruitment failed or was delayed in CK666-treated cells. Plo1 is recruited to the mitotic SPB just before the onset of mitosis by Pcp1, a pericentrin-related protein that localizes to the inner plaque of the SPB throughout the cell cycle. A *pcp1* mutant allele specifically incapable of recruiting Plo1 kinase to the SPB also shows defects in NE fenestration and SPB insertion failure [52]. This is in agreement with our observation that CK666 treated cells fail to form NE fenestration and show severe Plo1 kinase recruitment defects.

In addition to Plo1 kinase, Cut11 also failed to localize to the SPB. Cut11 is a seven-pass transmembrane protein, having homology to the Ndc1 protein in *S. cerevisiae*. Cut11 proteins are visible all over the nuclear envelope as it is associated with nuclear pore complex. However, during mitosis, it accumulates at the SPB in a punctuate manner from mitotic onset to mid-anaphase. The mutant *cut11.1* forms a fenestra but fails to insert the SPB and shows monopolar spindles emanating from a single SPB [37]. The fact that CK666 treated cells failed to recruit Cut11 to the SPB suggests that the defect is likely upstream of the SPB insertion event in mitosis.

The LINC (LInker of Nucleoskeleton and Cytoskeleton) complex consists of Sad1, an inner nuclear envelope SUN (Sad1-UNC-84) family protein, that associates with outer nuclear envelope KASH domain containing protein KMS1 and KMS2, and also tethers centromere via Csi1 in vegetative cells [40]. Sad1 reorganization into a ring-like pattern marks the beginning of NE fenestration [39]. This reorganization is significantly disturbed in CK666-treated cells, indicating severe impairment in NE fenestration. The defect in the reorganization of the Sad1 ring suggests that the upstream process that initiates Sad1 reorganization itself may be hampered when branched actin is disrupted. As it is challenging to capture NE fenestration in live cells with microscopy, we used the *cut12.1* strain expressing NLS-GFP as a proxy to understand this process. Cut12 is a protein needed for SPB insertion [16]. In the *cut12.1* mutant the NE fenestra forms but the duplicated SPB insertion is unsuccessful [16]. We exploited this fact to assess whether NE fenestration is impacted by branched actin disruption. Under restrictive conditions *cut12.1* mutants leak NLS-GFP through the fenestra. We do not see NLS-GFP leakage in CK666 treated *cut12.1* mutants further supporting that branched actin promotes NE fenestration. That CK666 does not impair mitotic entry in cells after pre-prophase release as with the tubulin mutant, *nda3-KM311*, further confirms that branched actin contributes to mitotic entry prior to SPB insertion.

The centromere-LINC complex is required for NE remodeling at the onset of mitosis, and defects in this contact compromise NE fenestration[17]. A few reports suggest the association of branched actin with the centrosome in metazoans, which supports chromatin organization and chromosome segregation and also acts as a perfusion barrier [23, 26, 27]. In the star fish oocytes, the nuclear envelope breakdown in mitosis is mediated by an F-actin shell formed by the Arp2/3 complex [43]. In light of these observations, there are two possibilities: either branched actin plays a direct role in NE remodeling, or its disruption alters LINC complex stability, thereby blocking NE fenestration. A close association of the Arp2/3 complex components such as Arc5 and actin bundling protein, Fim1 near the NE and the SPB supports a direct role for branched actin in fenestration.

While we see clear mitotic entry failure in CK666 treated cells, a fraction of these cells can enter mitosis albeit with delays. Further scrutiny of these cells shows severe mitotic progression abnormalities, as evidenced by irregular nuclear fission, multi-nucleated structures, a dissipated nuclear envelope, and severely bent late anaphase spindle microtubules. Careful analysis of these cells show that one of the SPBs fail to move away from the plane of the actomyosin ring and appears to be caught with it. This prevents one SPB from moving towards the cell end during anaphase and resulting in microtubule bending. In some cases, the SPB eventually escapes from this interaction while in others the association is not lost. This phenotype is completely lost in cells expressing LifeAct suggesting that the SPB likely associates with the actin in the actomyosin ring. LifeAct likely masks the actin site responsible for this interaction thus rescuing the phenotype. While this interaction may be an artefact of the lack of branched actin, it supports our observation that branched actin elements appear at the SPB in mitotic cells. Possibly, in the absence of branched actin, the SPB interacts with the only other available actin filament, namely the contractile ring. Future investigations will identify the component within the SPB that interacts with actin and its functional relevance.

While our findings strongly support a role for branched actin in NE fenestration, the mechanism by which this occurs is yet to be elucidated. Previous reports have shown that a functional Plo1 kinase is required for Sad1 ring formation and in mutants that fail fenestration, Plo1 does not accumulate at the SPB [39, 52]. We find that in CK666 cells, Plo1 accumulation to the SPB is either absent or significantly delayed. Thus, it is possible that branched actin promotes Plo1 accumulation which in turn enables NE fenestration. However, Plo1 accumulation to the SPB during mitosis depends on the cell-cycle dependent kinase, Cdc2. But in *cdc2* mutants Sad1 rings are able to form [36, 39]. This suggests that while functional Plo1 kinase is indeed required for Sad1 ring formation, its accumulation to the SPB at the onset of mitosis is likely not a requirement. Indeed, in *sad1* mutants where the ring does not form and fenestration fails, Plo1 levels are significantly reduced at the SPB. Thus, Plo1 accumulation at the SPB could be fenestration dependent. It is possible that the branched actin is required for Sad ring formation which then leads to fenestration. Another hypothesis is that the branched actin remodels the NE to make the fenestra, and when the SPB detects branched actin, it reorganizes Sad1 into a ring preparing for this event. While our report is the first to show the presence of branched actin components near the SPB at the onset of mitosis, Sad1 does bind the AP-1 adapter complex protein Apm1 [53]. Moreover, Cells lacking apm1 are temperature sensitive and show mitotic spindle separation defects[54]. Further investigation will determine if Apm1 at the SPB is related to the branched actin at this site.

The primary functions of branched actin include endocytosis, cell motility, cytokinetic abscission, and curvature sensing [24, 55, 56]. Branched actin’s role in mitosis is emerging in several eukaryotes, as evidenced by studies that project its association with chromosome congression, spindle assembly, and spindle size control[23, 26]. Branched actin has also been shown to promote nuclear envelope breakdown in starfish oocytes[29]. Our findings shed light on a novel function of the branched actin in mitotic entry and progression, plausibly by aiding NE fenestration. Further investigations are ongoing to understand how branched actin supports NE fenestration.

## Acknowledgements

We thank Sue Jaspersen, Megan King, Iain Hagan, Kathy Gould, David Kovar, James Moseley and Fred Chang for strains and Bret Judson for the Boston College imaging core facility.

## Materials and methods

### Strain and cell culture

The *S. pombe* strains used in this study are listed in Table S5. All strains are isogenic to the original strain PN567. The cells were maintained using YE culture medium at 25°C and grown exponentially for 48 hours before the experiments were performed. Standard techniques were used for genetic manipulation and analysis [57]

### Actin cytoskeleton disruptions

To block Arp2/3 complex mediated branched actin assembly, Cells were treated with 100 µM CK666 (SML006-5MG; Sigma Aldrich) in DMSO (D8418-250ML; Sigma-Aldrich) for 15 minutes. 10 µM Latrunculin (LatA; EMD Millipore) in DMSO was used to block the polymerization of all F-actin structures. For combinatorial treatment involving CK666+ LAT A, the same concentration of the drug as mentioned above was used. Control cells were treated with 0.1% DMSO in YE media.

### Workflow for the experiments involving genetic mutants

#### arp3-c1

*For experiments involving arp3-c1atb2-mCherry and arp3+ atb2-mCherry, cells* were grown at 32°C to mid-log phase and then shifted to restrictive temperature of 19°C for 12 hours. After 12 h. of incubation at 19°C, cells were shifted to semi permissive temperature of 22°C followed by imaging at one and two hours respectively using spinning disc confocal microscope. Respective temperature for imaging were maintained using the temperature chamber attached to the microscope.

#### cdc25-22

*cdc25-22* mutant strain was grown (in all the experiments) at 25°C to mid-log phase. Culture with an OD of 0.2 were shifted to 36°C for 4h followed by drug treatment (CK666/DMSO) for 15 minutes at 36°C. Cells were released to permissive temperature of 25°C and immediately proceeded with live-cell imaging at using spinning disc confocal microscope.

#### cut12.1

*cut12.1* and *cut12^+^* cells containing NLS-GFP cell was initially grown at 25°C to mid-log phase and shifted to 36°C with CK666/DMSO for 6hrs. Imaging was performed at regular intervals.

#### nda3-KM311

*nda3-KM311* strains were grown in YE medium at 25°C to mid-log phase. Then the cells were shifted to 18°C for 7hrs followed by treatment with CK666/DMSO for 15 min. Cells were released to permissive temperature of 34°C and Imaging were performed at 0 and 15 minutes of incubation at 34°C.

### Microscopy

A spinning disk confocal system that uses a Nikon Eclipse inverted microscope with a 100×1.49NA objective, a CSU-22 spinning disk system, and a Photometrics EM-CCD camera from Visitech International was used for live cell microscopy and for still images. Images were acquired using Metamorph (Molecular Devices) and analyzed using ImageJ/FIJI Bio-Formats plugins (National Institutes of Health).

We also used a 3i spinning disc confocal microscope with integrated Yokogawa spinning disk (Yokogawa CSU-X1 A1 spinning disk scanner) confocal on a Zeiss Axio Observer fully automated inverted microscope with a 100× −1.40 NA oil immersion objective and Teledyne Photometrics Prime 95b back-illuminated sCMOS camera (Serial No: A20D203014, Tucson, AZ). Images were acquired using SlideBook (3i Intelligent Imaging innovations, Denver, CO).

Super resolution microscopy was performed by a Zeiss LSM 880-Airyscan and a Plan-APOCHROMAT 63×, 1.4 NA oil objective using Zeiss Zen Black software. Images were processed using Airyscan processing mode in the Zeiss ZEN Blue 2.3 software. Structured illumination microscopy was performed using Zeiss Axio Observer.Z1 super-resolution microscope with Elyra S.1 system, that uses a Plan-Apochromat 63×/1.40 oil (DIC) objective. Images were acquired with a PCO-Tech Inc. pco.edge 4.2 sCMOS camera. The Zeiss Black software was used for channel alignment followed by structured illumination processing.

### Intensity measurements

Images acquired are further processed using FIJI image analysis tool. To measure the intensity of proteins, sum projected images was used after performing background subtraction using the plugin available in the FIJI. The region of interest was manually drawn at the respective region and the mean intensity was obtained.

To measure cdc25-nG nucleo-cytoplasmic ratios, the same ROI was used to measure the integrated density of both the nucleus as well as cytoplasm. N/C ratio for each cell, was calculated by dividing Nuclear integrated density values with the cytoplasmic integrated density.

### Structured illumination microscopy sample processing

For SIM, the cells were cultured at 25°C and shifted to 36°C for 4hrs followed by drug treatment and incubated at the same temperature for 15 min. Post incubation the cells were released to 25°C, followed by paraformaldehyde fixation at respective time points. For fixation, the cells were collected and treated with 32% formaldehyde (The final concentration was 4%) and incubated at room temperature for 10 min. Post incubation the cells were washed with ice cold PBS for 3min for 3 times and proceeded with slide preparation using prolong gold as a mountant. The slides were imaged after overnight incubation at room temperature.

### Statistics

All the experiments were performed in triplicates. Statistical analysis was performed using GraphPad Prism. A Student’s t test was used (two tailed, unequal variance) when comparing two conditions. One-way ANOVA followed turkey’s multiple comparison test, was used to determine significance for experiments with three or more conditions.

