## Supplementary Material for "Role of the Arp2/3 Complex in Regulating Mitotic Entry and Progression in *S. pombe*"

### Supplementary Figure 1

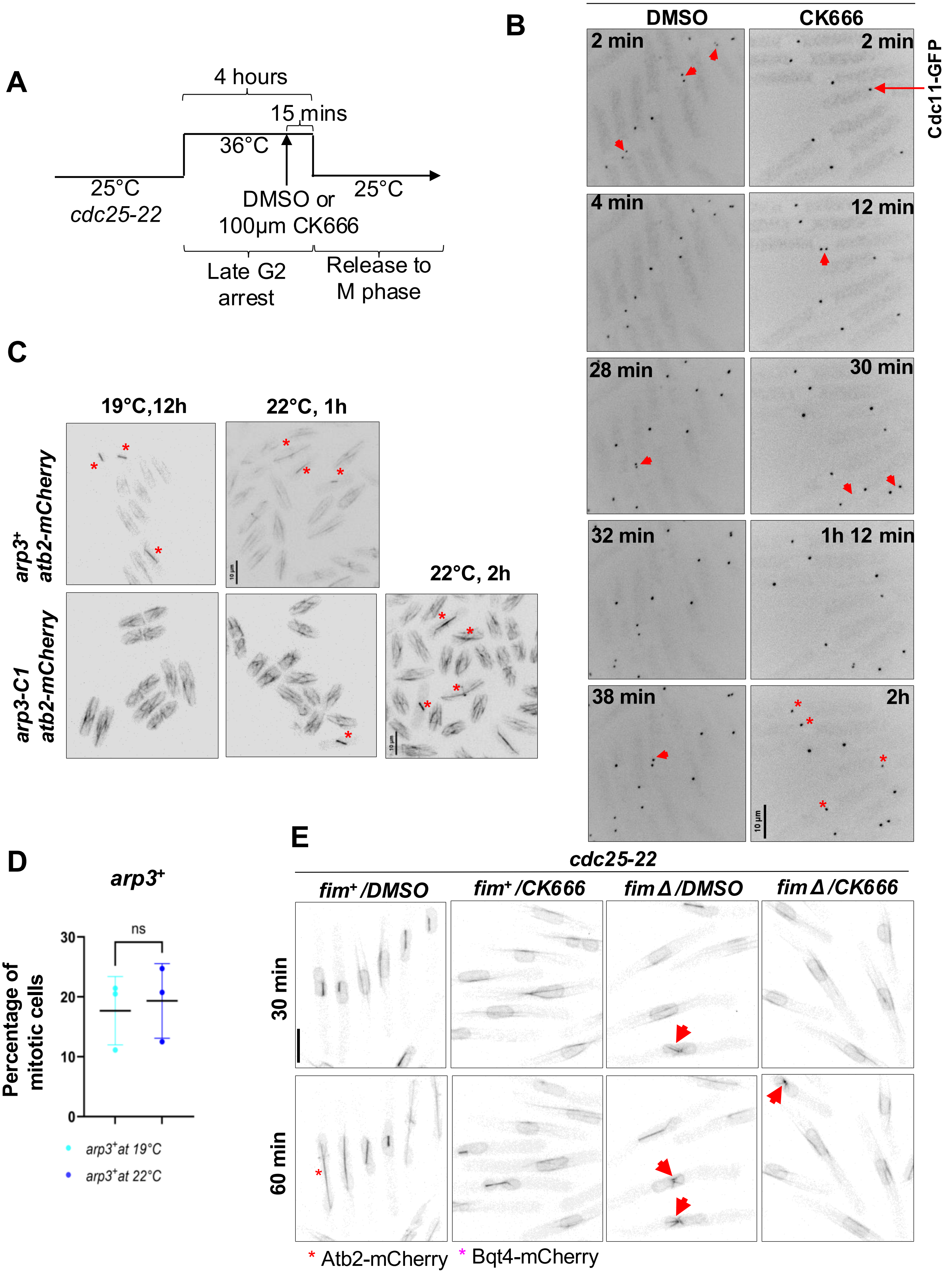

**Figure S1: Branched actin inhibition blocks or delays mitotic entry. A.** The graphical representation of workflow of *cdc25-22* cell-cycle arrest-release experiment. **B.** Time-lapse microscopy of DMSO or CK666-treated *cdc25-22* synchronized cells released from cell-cycle arrest. Sad1-mCherry marks SPB. Red arrowhead=cells entering mitosis. Red asterisk= cells failing mitosis. **C.** Mitotic index in *arp3*+ and *arp3-C1* at indicated temperatures and times. Atb2-mCherry labels microtubules. Red asterisk= mitotic cells. **D.** Graph shows the mitotic index of *arp3*+ cells at indicated temperatures. **E.** DMSO or CK666 treated synchronized *cdc25-22 fimΔ* cells 30 and 60 minutes after release from cell cycle arrest. bqt4-mCherry labels NE and atb2-mCherry labels microtubules. Red arrowhead shows monopolar spindles. Scale bar-10 μm.

#### Supplementary Figure 2

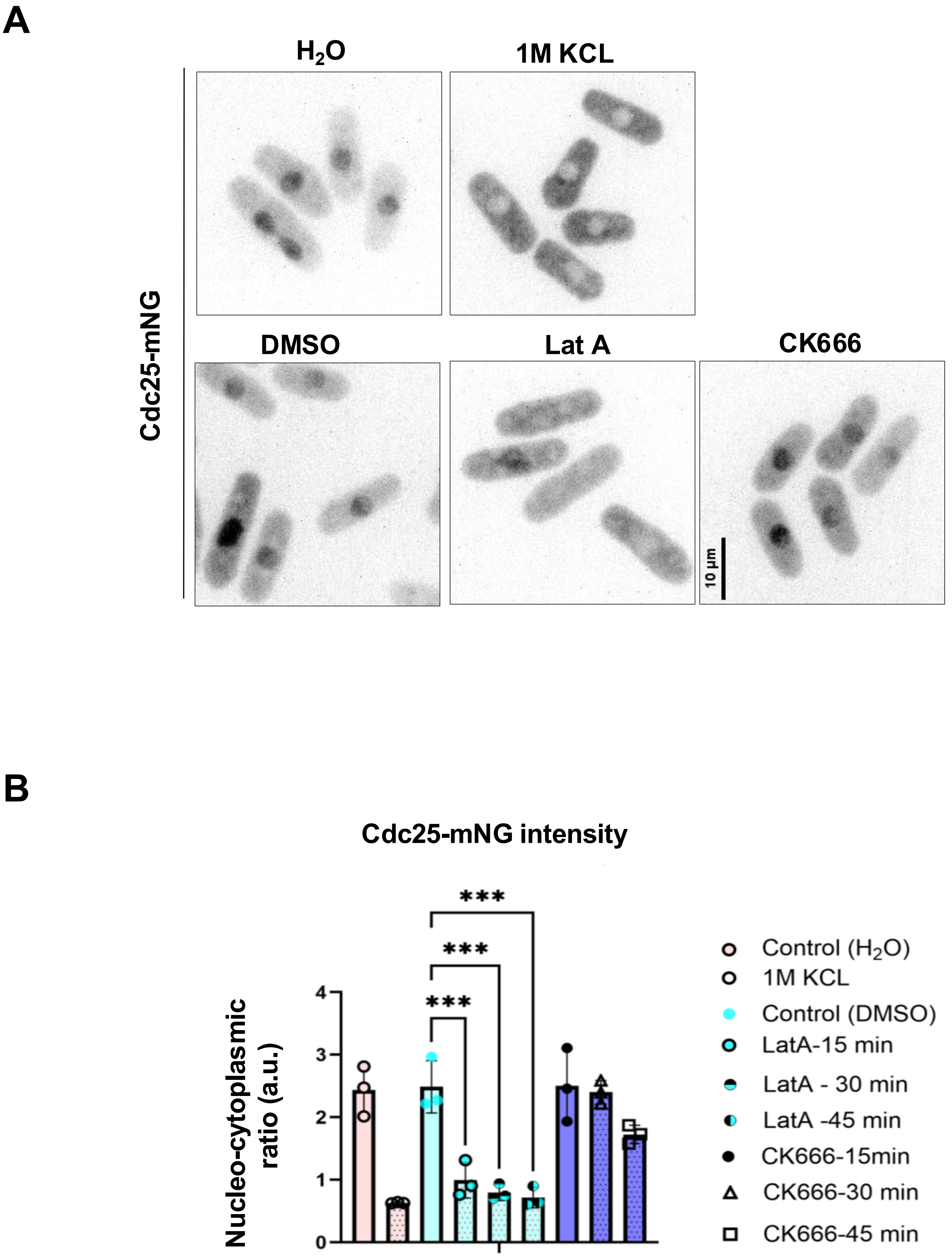

**Figure S2: CK666 treatment does not trigger a stress response. A.** Cdc25-mNG localization in cells treated as indicated. Scale bar-10 µm. **B.** Quantification of nucleocytoplasmic ratio of Cdc25-mNG in cells treated as indicated. n≥10, N=3, mean ± SD, one-way ANOVA, *** $p < 0.001$.

#### Supplementary Figure 3

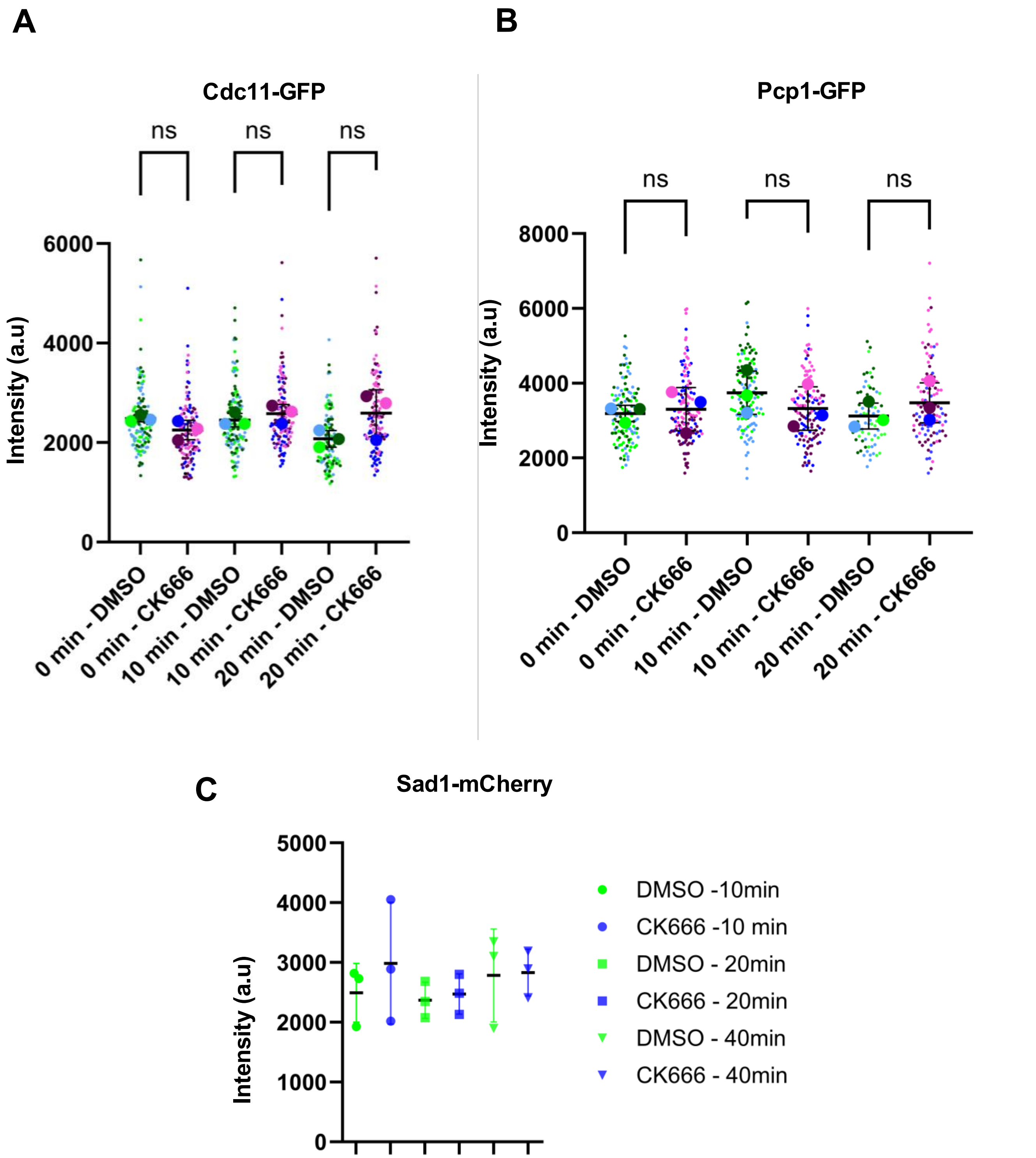

**Figure S3: loss of branched actin does not affect Cdc11, Pcp1 and Sad1 levels at the SPB.** Quantification of the **A.** Cdc11-GFP and **B.** PCP1-GFP and **C.** Sad1-mCherry intensities at the SPB in DMSO and CK666 treated *cdc25-22* synchronized cells released from cell cycle arrest for the indicated times. n≥22 cells, N=3, mean ± SD, One-way ANOVA, ns, not significant,

#### Supplementary Figure 4

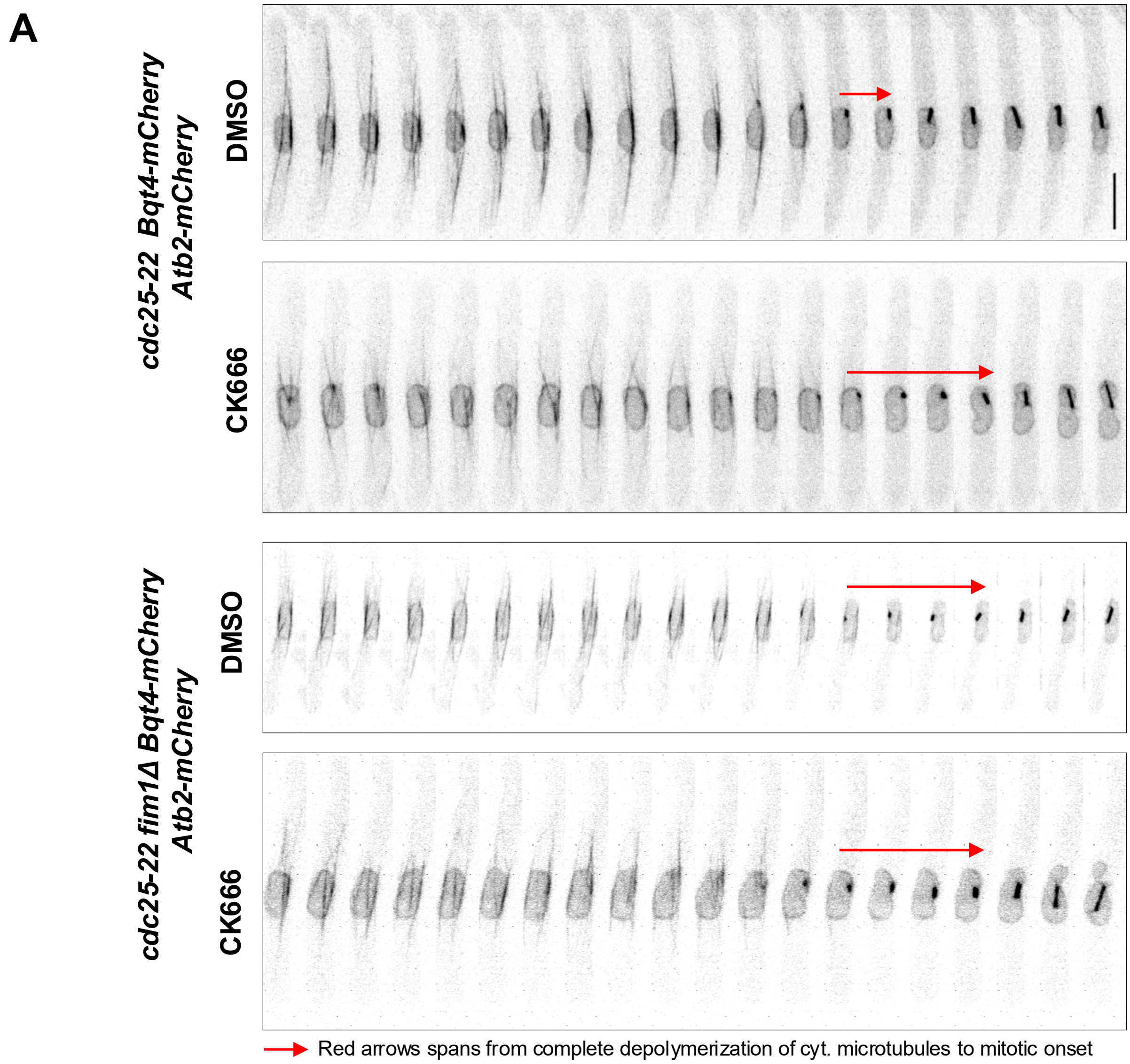

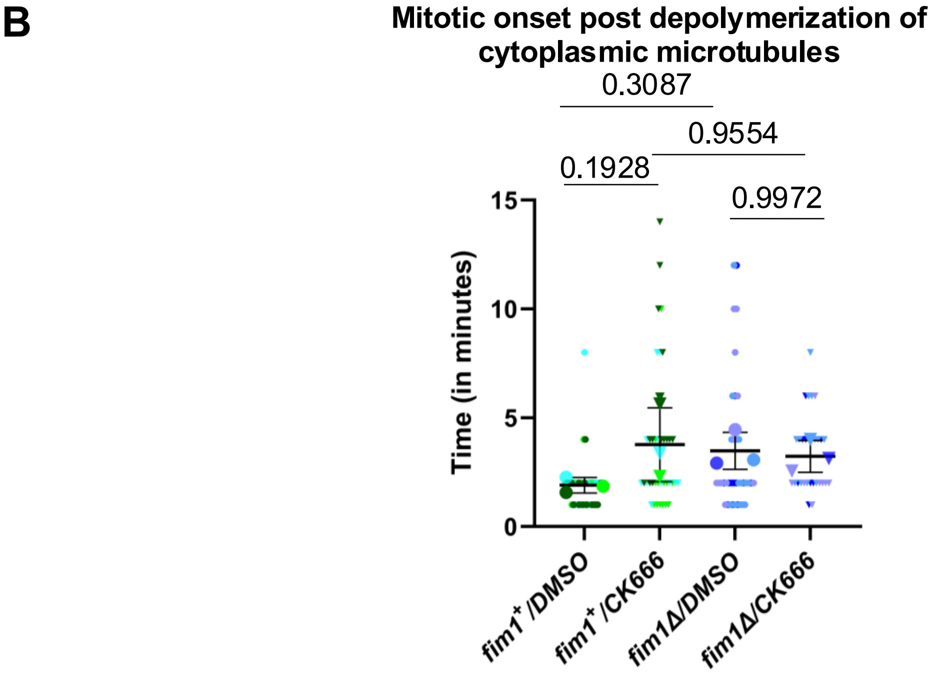

**Figure S4: Mitotic entry delay with respect to depolymerization of cytoplasmic microtubules. A.** Complete depolymerization of the cytoplasmic microtubules to mitotic onset in *fim1+* and *fim1Δ* cells treated with DMSO or CK666. **B.** Quantification of mitotic delay as shown above. p value, ns=not significant, one-way ANOVA, n≥15 cells, N=3, mean ± SD

#### Supplementary Fig 5

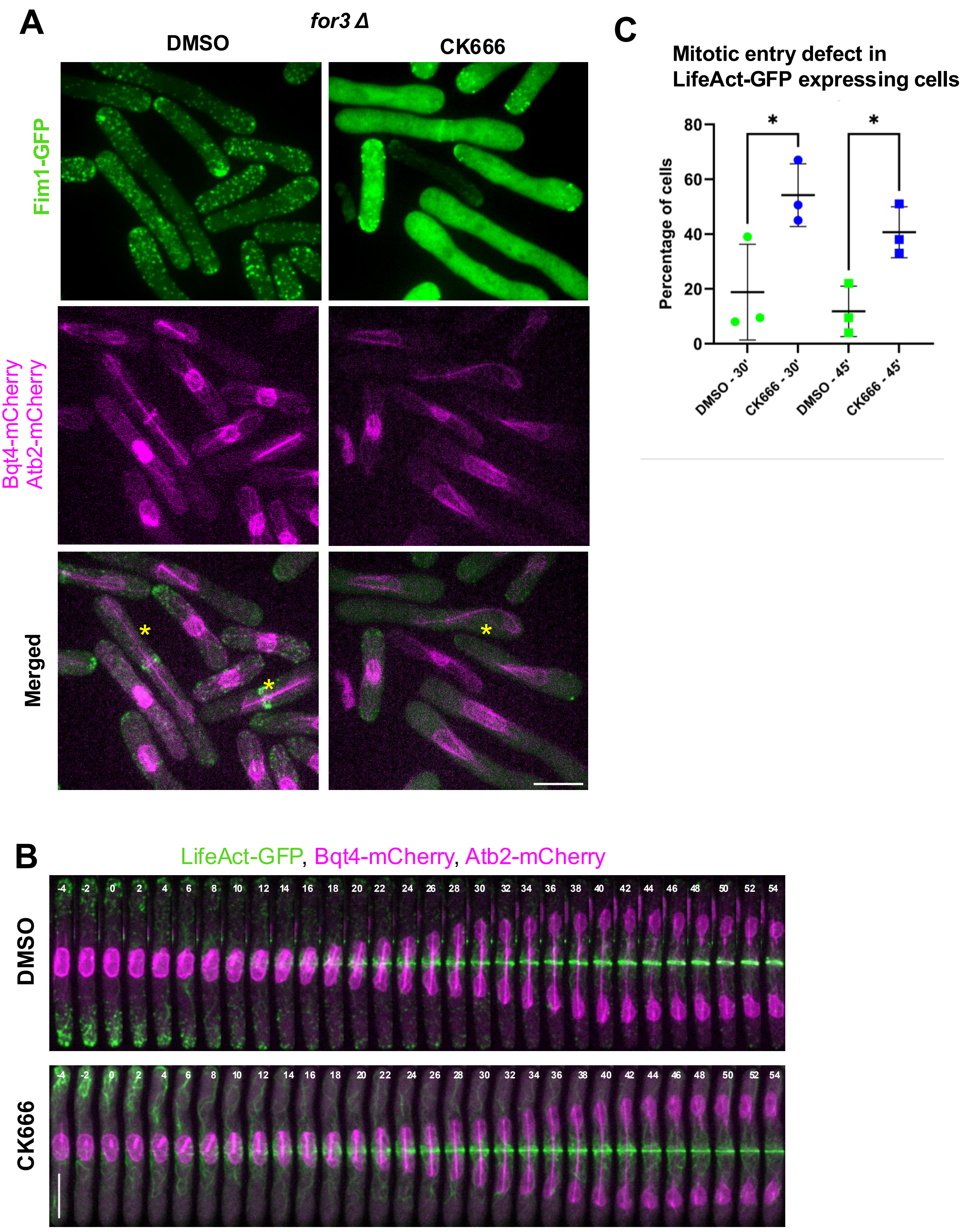

**Figure S5: Severe microtubule bending defects in the absence of branched actin require an intact actomyosin ring. A.** Spindle microtubule dynamics observed in DMSO or CK666 treated *cdc25-22 for3Δ* synchronized cells after release from cell-cycle arrest. Fim1-GFP label branched actin, Bqt4-mCherry labels NE and Atb2-mCherry labels spindle microtubules. **B.** Spindle microtubule dynamics during mitotic progression in DMSO or CK666 treated synchronized *cdc25-22* cells. Scale bar-10 µm. C. Mitotic entry defects in DMSO or CK666 treated *cdc25-22 LifeAct-GFP* synchronized cells after release from cell-cycle arrest. N≥21, N=3, mean ± SD, p value, * <0.05, one-way ANOVA.

#### Supplementary Videos

**Video S1:** Time-lapse microscopy of DMSO or CK666-treated *cdc25-22* synchronized cells released from cell-cycle arrest. Cdc11-GFP marks SPB.

**Video S2:** Plo1-GFP localization in DMSO or CK666-treated synchronous *cdc25-22* cells released into the cell cycle.

**Video S3:** Angular view (0-360°) of 3D reconstructed nucleus with Fim1-GFP localizing adjacent to Ppc89-mCherry 2 minutes prior to onset of mitosis.

**Video S4:** Angular view (0-360°) of 3D reconstructed nucleus with Fim1-GFP localizing to the nuclear envelope labeled with Bqt4-mCherry.

**Video S5:** Angular view (0-360°) of 3D reconstructed nucleus with Arc5-mCherry localizing to the nuclear envelope labeled with Bqt4-GFP.

**Video S6:** Nuclear fission in DMSO or CK666-treated synchronized *cdc25-22* cells post cell-cycle release. NE is labeled with Bqt4-GFP, SPB is labeled with Sad1-mCherry.

**Video S7:** Atb2-mCherry labeled spindle microtubule dynamics in DMSO or CK666-treated synchronized *cdc25-22* cells post cell-cycle release.

**Video S8:** SPB dynamics during mitosis relative to the actomyosin ring in DMSO or CK666-treated synchronized *cdc25-22* cells post cell-cycle release. SPB is labeled with Cdc11-GFP and the actomyosin ring is labeled with Rlc1-tdTomato.

#### Supplementary Tables

**Table S1:** Fim1-GFP localization at the SPB (Ppc89-mCherry) through a period of 10 minutes prior to SPB separation.

**Table S2:** Fim1-GFP localization at the nuclear envelope (Bqt4-mCherry) through a period of 18 to 8 mins prior to anaphase onset. Mitotic onset occurs 8 minutes prior to anaphase onset.

**Table S3:** Arc5-mCherry localization at the nuclear envelope (Bqt4-GFP) through a period of 18 to 8 minutes prior to onset of anaphase. Mitotic onset occurs 8 minutes prior to anaphase onset.

**Table S4:** Crn1-GFP localization at the SPB marked with Sad1-mCherry through a period of 12 minutes prior to SPB separation.

**Table S5:** Strain list

**Supplementary Table 1:** Fim1-GFP localization at the SPB (Ppc89-mCherry) through a period of 10 minutes prior to SPB separation.

| **Experiments** | **Mitotic cells** | **Time in Minutes (Prior to SPB separation)** | | | | | | | | | |
|---|---|---|---|---|---|---|---|---|---|---|---|
| | | **-10** | **-9** | **-8** | **-7** | **-6** | **-5** | **-4** | **-3** | **-2** | **-1** |
| Expt-1 | Cell 1 | x | x | x | ✓ | x | x | ✓ | x | x | x |
| | Cell 2 | x | x | x | x | x | x | x | x | ✓ | x |
| | Cell 3 | x | x | x | ✓ | x | x | x | x | ✓ | ✓ |
| | Cell 4 | ✓ | x | x | x | x | x | x | x | x | x |
| | Cell 5 | x | x | x | x | x | ✓ | x | x | x | x |
| | Cell 6 | x | x | x | x | x | x | ✓ | ✓ | x | x |
| | Cell 7 | x | x | ✓ | x | x | x | x | x | x | x |
| | Cell 8 | x | x | x | x | x | x | x | ✓ | x | ✓ |
| Expt-2 | Cell 1 | x | x | x | x | x | x | x | ✓ | x | x |
| | Cell 2 | x | x | x | x | x | x | x | ✓ | x | x |
| | Cell 3 | x | ✓ | x | x | x | x | x | x | x | x |
| | Cell 4 | x | x | x | x | x | ✓ | x | x | ✓ | x |
| | Cell 5 | x | x | x | ✓ | x | x | x | x | x | x |
| | Cell 6 | x | x | x | x | x | x | ✓ | x | x | x |
| | Cell 7 | x | x | x | ✓ | x | ✓ | x | x | x | ✓ |
| | Cell 8 | x | x | x | x | x | x | x | ✓ | x | ✓ |
| | Cell 9 | ✓ | x | x | x | x | x | x | x | x | ✓ |
| | Cell 10 | x | x | ✓ | x | x | x | x | x | x | x |
| | Cell 11 | x | x | ✓ | x | x | x | x | x | x | ✓ |
| | Cell 12 | x | x | x | x | x | x | ✓ | x | x | x |
| | Cell 13 | ✓ | x | ✓ | x | x | x | x | ✓ | ✓ | x |
| Expt-3 | Cell 1 | x | x | x | x | ✓ | x | x | x | x | x |
| | Cell 2 | x | ✓ | x | x | x | x | x | x | x | x |
| | Cell 3 | ✓ | x | x | x | x | x | x | x | x | x |
| | Cell 4 | x | x | x | x | ✓ | x | x | x | x | x |
| | Cell 5 | x | x | x | x | x | x | x | x | x | ✓ |
| | Cell 6 | x | ✓ | x | x | x | x | x | x | x | x |
| | Cell 7 | x | ✓ | x | x | x | x | x | x | x | x |
| | Cell 8 | x | x | x | x | x | x | x | ✓ | x | x |

**Supplementary Table 2**: Fim1-GFP localization at the nuclear envelope (Bqt4-mCherry) through a period of 18 to 8 mins prior to anaphase onset. Mitotic onset occurs 8 minutes prior to anaphase onset.

| Experiments | Mitotic cells | | Time in Minutes (Prior to Anaphase Onset) | | | | | | | | | |
|---|---|---|---|---|---|---|---|---|---|---|---|---|
| | | -18 | -17 | -16 | -15 | -14 | -13 | -12 | -11 | -10 | -9 | -8 |
| Expt-1 | Cell 1 | x | ✓ | ✓ | x | ✓ | ✓ | ✓ | ✓ | ✓ | ✓ | ✓ |
| | Cell 2 | x | x | ✓ | x | ✓ | | ✓ | x | ✓ | ✓ | ✓ |
| | Cell 3 | x | x | ✓ | x | ✓ | ✓ | x | ✓ | ✓ | ✓ | ✓ |
| | Cell 4 | x | ✓ | ✓ | ✓ | x | x | ✓ | ✓ | ✓ | ✓ | ✓ |
| | Cell 5 | x | x | ✓ | x | x | x | x | x | ✓ | ✓ | ✓ |
| Expt-2 | Cell 1 | x | ✓ | ✓ | ✓ | ✓ | ✓ | ✓ | ✓ | ✓ | ✓ | ✓ |
| | Cell 2 | ✓ | ✓ | ✓ | x | ✓ | ✓ | ✓ | x | x | x | x |
| | Cell 3 | ✓ | x | x | x | x | ✓ | ✓ | x | x | ✓ | x |
| | Cell 4 | ✓ | x | x | x | x | x | x | ✓ | x | ✓ | ✓ |
| | Cell 5 | x | x | x | ✓ | ✓ | ✓ | x | ✓ | x | x | x |
| Expt-3 | Cell 1 | ✓ | x | x | x | x | x | x | x | ✓ | ✓ | ✓ |
| | Cell 2 | x | ✓ | x | x | x | x | x | x | ✓ | ✓ | ✓ |
| | Cell 3 | x | x | ✓ | ✓ | x | x | x | x | ✓ | ✓ | ✓ |
| | Cell 4 | x | x | x | x | ✓ | x | ✓ | ✓ | ✓ | ✓ | ✓ |
| | Cell 5 | x | ✓ | x | x | ✓ | ✓ | ✓ | x | x | ✓ | ✓ |

**Supplementary Table 3**: Arc5-mCherry localization at the nuclear envelope (Bqt4-GFP) through a period of 18 to 8 minutes prior to onset of anaphase. Mitotic onset occurs 8 minutes prior to anaphase onset.

| Experiments | Mitotic cells | Time in Minutes (Prior to Early Anaphase) | | | | | |
|---|---|---|---|---|---|---|---|
| | | -18 | -16 | -14 | -12 | -10 | -8 |
| Expt-1 | Cell 1 | x | x | ✓ | ✓ | x | ✓ |
| | Cell 2 | x | ✓ | ✓ | x | ✓ | ✓ |
| | Cell 3 | ✓ | ✓ | x | x | x | ✓ |
| | Cell 4 | ✓ | ✓ | ✓ | ✓ | ✓ | ✓ |
| Expt-2 | Cell 1 | x | ✓ | ✓ | x | ✓ | ✓ |
| | Cell 2 | ✓ | x | ✓ | x | x | ✓ |
| | Cell 3 | x | ✓ | ✓ | ✓ | ✓ | ✓ |
| | Cell 4 | x | ✓ | ✓ | ✓ | x | x |
| Expt-3 | Cell 1 | ✓ | x | x | ✓ | ✓ | x |
| | Cell 2 | x | ✓ | ✓ | ✓ | x | x |

**Supplementary Table 4**: Crn1-GFP localization at the SPB marked with Sad1-mCherry through a period of 12 minutes prior to SPB separation.

| **Experiments** | **Mitotic cells** | **Time in Minutes (Prior to SPB separation)** | | | | | | | | | | | |
|---|---|---|---|---|---|---|---|---|---|---|---|---|---|
| | | **-12** | **-11** | **-10** | **-9** | **-8** | **-7** | **-6** | **-5** | **-4** | **-3** | **-2** | **-1** |
| Expt-1 | **Cell 1** | x | x | x | x | x | x | x | x | x | x | x | x |
| | Cell 2 | ? | x | ✓ | x | x | x | x | x | ✓ | x | x | x |
| | Cell 3 | ? | x | x | ✓ | x | x | x | x | x | ✓ | x | ✓ |
| | Cell 4 | ✓ | x | x | x | x | x | ✓ | x | x | ✓ | x | ✓ |
| | Cell 5 | ✓ | ✓ | x | x | x | x | x | ✓ | x | ✓ | x | x |
| | Cell 6 | ✓ | ✓ | x | x | ✓ | ✓ | x | x | x | x | x | |
| | Cell 7 | x | x | x | x | x | x | x | x | x | x | x | x |
| | Cell 8 | x | x | x | x | x | x | x | x | x | x | x | x |
| Expt-2 | Cell 1 | x | ✓ | x | x | x | x | x | x | x | x | x | x |
| | Cell 2 | x | x | x | x | x | x | x | x | x | x | x | x |
| | Cell 3 | x | x | x | x | x | ✓ | x | x | x | x | ✓ | ✓ |
| | Cell 4 | x | x | x | x | ✓ | x | x | x | x | ✓ | x | x |
| | Cell 5 | x | x | ✓ | x | x | x | x | x | ✓ | x | ✓ | x |
| Expt-3 | Cell 1 | ✓ | x | x | ✓ | ✓ | ✓ | ✓ | x | x | x | ✓ | ✓ |
| | Cell 2 | x | x | ✓ | ✓ | x | x | x | x | x | x | x | x |
| | Cell 3 | ✓ | x | ✓ | x | ✓ | x | ✓ | x | x | ✓ | ✓ | x |
| | Cell 4 | x | x | x | x | x | x | x | ✓ | x | x | x | x |
| | Cell 5 | x | x | ✓ | x | ✓ | x | x | ✓ | ✓ | x | x | x |
| | Cell 6 | x | x | ✓ | x | x | x | ✓ | x | ✓ | ✓ | x | x |
| | Cell 7 | x | x | x | x | x | x | x | x | x | x | x | x |
| | Cell 8 | x | x | x | x | x | x | x | x | x | x | x | x |
| | Cell 9 | x | x | x | x | x | ✓ | x | x | x | ✓ | x | x |
| | Cell 10 | x | x | x | x | x | x | ✓ | x | x | x | x | x |

**Table S5: Strain list.**

| ***Strain*** | ***Genotype*** | ***Source*** |
|---|---|---|
| *YMD1956* | *cdc25-22 cdc11-GFP: KanMX6 sad1-mCherry: KanMX ade6 leu1-32 ura4-D18* | *This Study* |
| *YMD1990* | *cdc25-22 sad1-mCherry: KanMX6 bqt4-GFP: aur1R leu1-32 ura4-D18 his3-D1 ade6* | *This Study* |
| *KGY11659* | *cdc25-22 plo1-GFP3: KanMX6 sad1-mCherry: KanMX ade6-M210 leu1-32 ura4-D18* | *Kathy Gould* |
| *YMD1999* | *cdc25-22 atb2-mCherry: natMX ade6 Leu1-32 ura4-D18* | *This Study* |
| *JM 5290* | *cdc25-Neon Green* | *James Mosely* |
| *YMD2411* | *cdc25.22 atb2-mCherry: NatMX bqt4-mCherry: aur1R LifeAct-GFP:Leu+ ura4-D18 his3-D1 ade6* | *This Study* |
| *YMD2435* | *cdc25-22 for3Δ: KanMX6 atb2-mCherry: NatMX bqt4-mcherry: aur1R fim1-GFP: KanMX6 ura4-D18 ade6 leu1-32* | *This Study* |
| *YMD2330* | *cdc25-22 atb2-mCherry: NatMX bqt4-mcherry: aur1R fim1-GFP: KanMX6 ura4-D18 ade6 leu1-32* | *This Study* |
| *YMD2486* | *cdc25.22 fimΔ: KanMX6 bqt4-GFP: aur1R atb2- mCherry:NatMX ade6 leu1-32 ura4-D18* | *This Study* |
| *YMD2329* | *cdc25.22 bqt4-GFP: aur1R atb2- mCherry:NatMX ade6 leu1-32 ura4-D18* | *This Study* |
| *YMD2137* | *cdc25-22 cut11-GFP: ura4+ sad1-mCherry: KanMX6 ade6 leu1-32 Ura4-D18* | *This Study* |
| *YMD2636* | *cdc25-22 cdc11-GFP: KanMX6 rlc1- tdTomato:NatMX ade6 leu1-32 ura4-D18* | *This Study* |
| *YMD2674* | *fim1-GFP: KanMX6 Ppc89-mCh: NatMx6 ade6 leu1-32 ura4-D18* | *This Study* |
| *YMD2166* | *arc5-mCherry: NatMX4 bqt4-GFP: aur1R leu1-32 ura4-D18 his3-D1 ade6-210* | *This Study* |
| *YMD2038* | *fim1-GFP: KanMX6 bqt4-mCherry: aur1R leu1-32 ura4-D18 his3-D1 ade6-210* | *This Study* |
| *YMD2363* | *arp3-c1 atb2-mCherry: NatMX* | *This Study* |
| *YMD 1964* | *bqt4-GFP: aur1R sad1-mCherry: KanMX6 leu1-32 ura4-D18 his3-D1 ade6-210* | *This Study* |
| *YMD 2000* | *nda3.1 sad1-mCherry: KanMx* | *This Study* |
| *1477* | *pcp1-GFP: hphMX6 plo1-mCherry: KanMX6 ura4-D18 leu1-32* | *Megan King* |
| *1015* | *cut11-GFP: ura4+ sad1-mCherry: hphMX6 h+ ura4-D18 leu1-32 lys1?* | *Megan King* |
| *fySLJ985* | *pREP3X-SV40 NLS-GFP-lacZ, Ppc89-mCherry-NatMX6, cut12.1, leu1-32, ura4-D18* | *Sue Jaspersen* |
| *KV609* | *h+, fim1Δ: kanMX6, lifeact-GFP:leu+, ade6-m216, leu1-32, ura4-D18.* | *David Kovar* |
| *AY361* | *arc5-mCherry-NatMX4 ade6-M216 leu1-32 ura4-D18* | *Fred Chang* |
